# Post-fluoroquinolone treatment molecular events and nutrient availability modulate *Staphylococcus aureus* antibiotic persistence

**DOI:** 10.1101/2025.06.26.661800

**Authors:** Jonathan I. Batchelder, Nisha Mahey, Wendy W. K. Mok

## Abstract

*Staphylococcus aureus* is a bacterial pathogen associated with about one million deaths per year. *S. aureus* infects diverse host sites including skin and the airway. At nutrient-limited infection sites, *S. aureus* cells become increasingly metabolically quiescent. While lower metabolic activity may help contain growth of the pathogen, it can enhance *S. aureus*’s persistence to antibiotic treatment. Here, we focus on starved *S. aureus*’s response to fluoroquinolones (FQs), which inhibit topoisomerases necessary for DNA and RNA synthesis and lead to double-stranded DNA break (DSB) formation. We show that although FQs accumulate in stationary-phase *S. aureus* during treatment, DSBs form and cells start to die during the post-treatment period when nutrients are replenished. We found that most persisters suffer DNA damage and rely on RecA to repair DSBs post-treatment. We discovered that persisters frequently pass on damaged chromosomes to their progeny, causing many of their progeny to stall. Given that persister resuscitation and death of non-persisters occur during the post-FQ treatment period, we asked how nutrient availability during this critical time impacts *S. aureus* survival. We show that continued starvation after treatment ends enhances *S. aureus* survival, even in populations lacking DSB-repair mechanisms. We demonstrate that post-treatment starvation delays the resumption of nucleic acid synthesis, presumably limiting topoisomerase activity, and allows the cells time to remove some intracellular FQs before they can trap active topoisomerases. Collectively, our findings highlight DSB repair processes and environmental conditions that can be targeted to improve treatment outcomes for staphylococcal infections.

**Significance Statement:** *S. aureus*’s ability to overcome antibiotic treatment makes it a formidable pathogen. Since nutrients are often limited at infection sites, understanding how starved *S. aureus* responds to clinically relevant antibiotics, including DNA-damaging FQs, is essential for improving treatment outcomes. We show that DNA damage accumulates during the post-treatment period when nutrients are replenished. Continuing to starve the cells after treatment limits the incidence of DSBs, helping some cells survive even if they are incapable of DSB repair. Finally, we show that offspring stemming from cells that survive FQ treatment often experience damage and stress for several generations. Our work highlights strategies *S. aureus* deploys to recover from FQ treatment that can be targeted to enhance antibiotic efficacy.

## Introduction

*Staphylococcus aureus* is a difficult-to-treat opportunistic pathogen that infects a plethora of human host sites, including the bloodstream, respiratory tract, endocardium, skin, and bone (1). To successfully colonize and chronically infect such diverse sites, *S. aureus* must adapt to non-ideal nutrient sources to overcome host nutrient sequestration, which limits the availability of metal ions and metabolites like glucose, *S. aureus*’s preferred carbon source (2–7). Nutrient limitation in experimental cultures and at infection sites has been shown to reprogram bacterial metabolism and help *S. aureus* survive antibiotic treatment (8–13). Specifically, decreased nutrient availability and metabolic activity increase the proportion of bacterial persisters—phenotypic variants in clonal cultures that can survive antibiotic treatment regimens that kill their kin. As persisters tolerate antibiotics and resume growth, they can reseed an infection after antibiotic treatment ends. Further exacerbating the problem, persisters have also been shown to promote resistance development (14–17).

Although starved *S. aureus* cultures are tolerant to many antibiotics, particularly those that inhibit cell wall synthesis, fluoroquinolones (FQs) retain some bactericidal activity against these recalcitrant populations (8–11, 18–20). Among these drugs, ciprofloxacin (CIP) is approved for treating *S. aureus* skin and skin structure infections (SSSIs), and delafloxacin (DLX), which gained FDA approval in 2017, is indicated for treating staphylococcal SSSIs as well as respiratory infections (21, 22). FQs bind and inhibit the catalytic activity of topoisomerases, which are required for both DNA and RNA synthesis, and are thought to cause double-stranded DNA breaks (DSBs) when the FQ-topoisomerase-nicked DNA complexes become irreversibly trapped (23–26). Understanding how *S. aureus* persisters survive FQ treatment is essential for improving pathogen clearance and patient outcomes following antibiotic therapy.

Building on the strong evidence that reduced metabolic activity enhances antibiotic tolerance/persistence, several groups, including ours, have demonstrated that stimulating the metabolism of stationary-phase (starved) bacteria during antibiotic treatment decreases survival to multiple drug classes (27–32). Our group and others previously demonstrated that adding nutrients to stimulate stationary-phase cultures of *Escherichia coli* and *S. aureus* reduces persistence to fluoroquinolones (FQs) (10, 11, 20, 27). Metabolic stimulation decreased *E. coli* persistence by enhancing RNA synthesis and requisite topoisomerase activity, whereas it decreased *S. aureus* persistence mainly by increasing oxidative stress (20, 27). These mechanistic differences emphasize that although *E. coli* paradigms are helpful starting points for formulating hypotheses, there are nuances in how individual bacterial species respond to antibiotics; therefore, different strategies are required to eliminate persisters from distinct pathogens.

Beyond the effects of nutrient availability during FQ treatment, several studies have demonstrated the importance of *recA, recB*, and other genes involved in DNA recombination and DSB repair for *E. coli* cells stemming from stationary-phase cultures to survive FQ treatment (33– 38). Loss of *recA* or *recB* reduces *E. coli* FQ persistence by ∼1,000-fold, suggesting that most FQ persisters rely on DSB repair mechanisms to overcome the effects of FQ treatment rather than simply avoiding antibiotic-induced damage (38). In *E. coli* and other species, it has been demonstrated that prolonged nutrient limitation that perpetuates after FQ treatment is over can enhance persistence (35, 38–40). In particular, post-treatment starvation enhances *E. coli* persistence by allowing cells to repair DSBs via RecA before nutrient replenishment causes metabolic and biosynthetic processes to resume (38). The impact of starvation following FQ treatment on the survival of *S. aureus* has not been established.

In this study, we demonstrate that *S. aureus* persisters, like *E. coli*, engage DSB-repair mechanisms after FQ treatment ends, and post-treatment starvation increases *S. aureus* persistence. We also uncovered important distinctions between how *S. aureus* and *E. coli* persisters survive FQ treatment. Unlike *E. coli*, post-treatment starvation increases *S. aureus* persistence even in cells that cannot repair DSBs. We show that starvation delays the resumption of nucleic acid synthesis so that intracellular FQ abundance can decrease before topoisomerases reactivate. When we tracked persister outgrowth, we found that *S. aureus* persisters can pass DNA damage to their progeny, leading to stress that spans multiple generations. This is unlike *E. coli* persisters, which appear to fully repair FQ-induced DNA damage before producing healthy progeny (39). In all, our findings offer substantial insights concerning the mechanisms of persistence in the clinically significant pathogen *S. aureus*, which could inform future strategies for improving treatment outcomes.

## Results

### Loss of *recA* decreases FQ persistence of stationary-phase *S. aureus*

In this study, we used the methicillin- and FQ-sensitive *S. aureus* strain HG003 to explore how the nutrient environment during and after FQ treatment impacts FQ persistence (41). We hypothesize that *S. aureus* FQ persisters from stationary-phase cultures suffer DNA damage that they must repair to survive. To test this hypothesis, we assessed the impact of disrupting *recA* on *S. aureus* survival following FQ treatment. We confirmed that both wild-type (WT) HG003 and its *recA*::Tn derivative are in stationary phase after 18 h of growth **(Fig. S1A-B)**. Then, we treated both these strains with DLX and CIP, at a range of doses above their minimum inhibitory concentrations (MICs), for 5 h. As expected, disruption of *recA* and subsequent loss of HR capabilities significantly reduced survival by ∼100-fold across the range of DLX and CIP concentrations tested, as shown previously for nalidixic acid with growing cultures of *E. coli* **(Fig. 1A-B)** (42).

**Figure 1.**
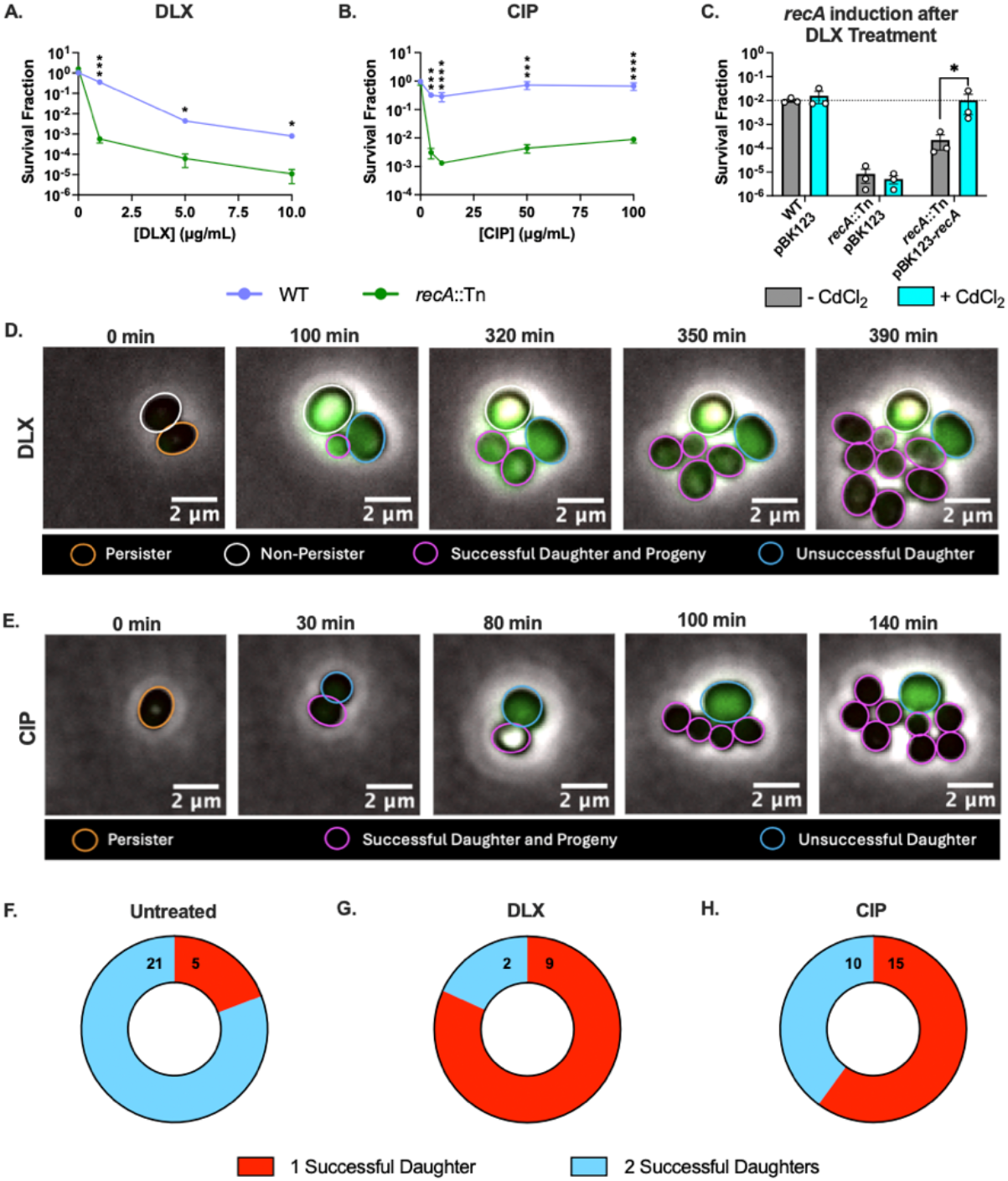
RecA is required during post-antibiotic treatment recovery for maximal FQ persistence. (A-B) Stationary-phase cultures of *S. aureus* WT or *recA*::Tn were treated with designated doses of (A) DLX or (B) CIP for 5 h. (C) WT with pBK123, *recA*::Tn with pBK123, and *recA*::Tn bearing pBK123-*P*_*cad*_-*recA* were treated with 5 μg/mL DLX for 5 h before recovery on media with or without CdCl_2_. Dotted line indicates survival fraction of DLX-treated WT populations recovered without CdCl_2_. (D-E) HG003::*P*_*recA*_*-gfp* was monitored via timelapse microscopy after treatment with (D) DLX or (E) CIP. Green cells are positive for *gfp* expression (indicating *recA* induction), and red cells are positive for PI staining (indicating loss of membrane integrity in dying/dead cells). (D) The DLX non-persister (white outline) expresses *recA* and does not divide throughout 16 h of imaging. The DLX persister (orange outline) swells then divides. The successful daughter (magenta outline) divides, giving rise to more progeny that express *recA*, but the unsuccessful daughter (blue outline) stalls and expresses *recA*. (E) The CIP persister (orange outline) divides into two cells. The unsuccessful daughter (blue outline) expresses *recA* and swells but never divides. The successful daughter (magenta outline) divides and gives rise to more progeny. (F-H) The proportion of persisters producing one or two successful daughter cells (daughters that produce progeny rather than stalling or dying) after treatment with (F) DMSO (untreated control), (G) 5 μg/mL DLX, or (H) 10 μg/mL CIP was calculated. At least three independent replicates were performed for each survival experiment, and microscopy images are representative of at least two independent replicates. (A-C) *P* values were calculated using *t*-tests assuming (A-B) unequal variances to compare the log-transformed survival fractions at each dose or (C) equal variances to compare the log-transformed survival fractions between cultures recovered with or without CdCl_2_. \**P* < 0.05, \*\*\**P* < 0.005, \*\*\*\**P* < 0.001. Only statistically significant comparisons are shown. Error bars denote SEM.

Complementing *recA* into the genome of *recA*::Tn restored survival to WT levels, confirming that loss of *recA* is responsible for increased killing by FQs in stationary-phase *S. aureus* **(Fig. S2A-B)**. For subsequent experiments, we used 5 μg/mL DLX (∼2500x WT MIC) and 10 μg/mL CIP (∼333x WT MIC). We chose these doses because the survival of WT populations had plateaued at these concentrations and the differences in survival between WT and *recA*::Tn were significant. Additionally, they reflect physiological concentrations achieved during clinical administration **(Fig. 1A-B)** (43, 44). When we sampled *S. aureus* treated with these DLX and CIP doses at different times, we detected biphasic killing within 5 h, indicating that the surviving cells were persisters within a phenotypically heterogeneous culture **(Fig. S3A-B)**(16).

### *S. aureus* expresses *recA* during the post-treatment recovery period

After establishing the importance of *recA* for stationary-phase *S. aureus* surviving FQ treatment, we asked whether inducing *recA* during the post-FQ treatment recovery period is sufficient to rescue survival. To answer this question, we complemented a cadmium-inducible copy of *recA* into *recA*::Tn (45). We found that the survival of DLX-treated *recA*::Tn was comparable to that of WT cells even when *recA* was induced only during the post-treatment period **(Fig. 1C)**. These results suggest that *S. aureus* persisters mainly repair FQ-induced DNA damage once treatment terminates and nutrients are replenished, rather than during FQ treatment itself.

To further gauge when FQ-treated *S. aureus* cells induce *recA* expression, we used timelapse fluorescence microscopy to track *S. aureus* harboring a plasmid-borne *recA* transcriptional reporter following treatment with DLX or CIP. We found that stationary-phase *S. aureus* persisters and non-persisters both maintained membrane integrity and did not take up propidium iodide (PI) throughout treatment. Both persisters and cells that die expressed *recA* during the post-treatment recovery period, but the treatment-free control remained dim **(Fig. 1D-E; Fig. S4-5; Movies S1-6)**. The expression of *recA* during recovery suggests that both persisters and their dying counterparts attempt to repair FQ-induced DSBs via RecA-mediated processes after treatment ends. This finding implies that many *S. aureus* FQ persisters do not survive simply by escaping antibiotic-induced damage.

Beyond *recA* expression in persisters, we tracked the fates of persister progeny. As expected, most untreated cells (21/26; 81%) divided and gave rise to two daughter cells that both successfully produced progeny **(Fig. 1F; Fig. S5B-C; Movies S5-6)**. However, DLX-treated persisters rarely produced two successful daughter cells (2/11 persisters; 18%), and only 40% (10/25) of CIP-treated persisters produced two successful daughters **(Fig. 1G-H)**. Instead, we frequently observed that one of a persister’s daughters proliferated to form a microcolony while the other daughter failed to divide further and sometimes died. The unsuccessful daughter often continued to induce P_*recA*_*-gfp* after separating from the successful daughter **(Fig. 1D-E; Fig. S5D-E; Movies S1-4)**. To confirm that this observation was not an artefact of our *recA* reporter system, we repeated these experiments with a reporter-free strain and found that its *S. aureus* FQ persisters likewise frequently produce only one successful daughter **(Fig. S6A-D; Movies S7-10)**. These findings suggest that while *S. aureus* persisters engage RecA and attempt to repair FQ-induced DNA damage during the post-FQ recovery period, some damage may not be completely repaired before the persister divides.

### Both persisters and their progeny appear to experience damage from FQ treatment

Given our finding that one of a persister’s daughters often shows high *recA* expression and fails to divide, we hypothesized that this unsuccessful daughter inherits a damaged chromosome from the persister while the successful daughter inherits an intact chromosome. This is a reasonable assumption since a recent report showed that most stationary-phase *S. aureus* cells contain two chromosomes (46). To test this hypothesis, we integrated a RecA-GFP translational fusion into the chromosome of *S. aureus* HG003. This strategy has been used in *E. coli* and *S. enterica*, resulting in DSBs being marked with green foci as RecA-GFP binds to initiate repair (**Fig. S7A**) (40, 47). We confirmed that untreated cells bearing this reporter divide normally and do not usually produce foci before dividing **(Fig. S7B-F)**. In the absence of FQ treatment, a cell at the start of imaging [Generation 0 (G0)] usually divided into two cells at G1. Then, each G1 cell divided into two more cells, forming four cells at G2, which in turn gave rise to eight cells at G3 **(Fig. S7C, E)**. Later generations sometimes formed faint foci, likely due to transient DSBs that form and are repaired during normal DNA replication and division **(Movies S11-12)** (48). On the other hand, most FQ-treated non-persisters (dying cells) had bright foci and never divided, while a small fraction divided once before both daughters stalled **(Fig. S8; Movies S13-16)**.

Next, we tracked the formation and inheritance of foci in DLX and CIP persisters **(Fig. 2A-D; Fig. S9)**. If persisters partition an intact chromosome to one daughter and a damaged chromosome to the other, we would expect to see the persister form a focus before dividing then pass this focus on to one of its daughters. Indeed, unlike untreated cells, most DLX (14/18; 78%) and CIP (14/17; 82%) persisters formed a focus before dividing **(Fig. 2E-F;** compare to Fig. S7F). Occasionally, only one of a persister’s two daughters inherited a focus and stalled, and the focus-free daughter continued to divide **(Fig. 2A-B)**. However, 50% of DLX persisters (9/18) and a large proportion of CIP persisters (15/17; 88%) produced two daughters with foci **(Fig. 2C-D, G-H; Fig. S9A-B)**. This observation suggests that, contrary to our hypothesis, both daughters of an *S. aureus* FQ persister may frequently have DSBs that they must repair if they are to survive.

**Figure 2.**
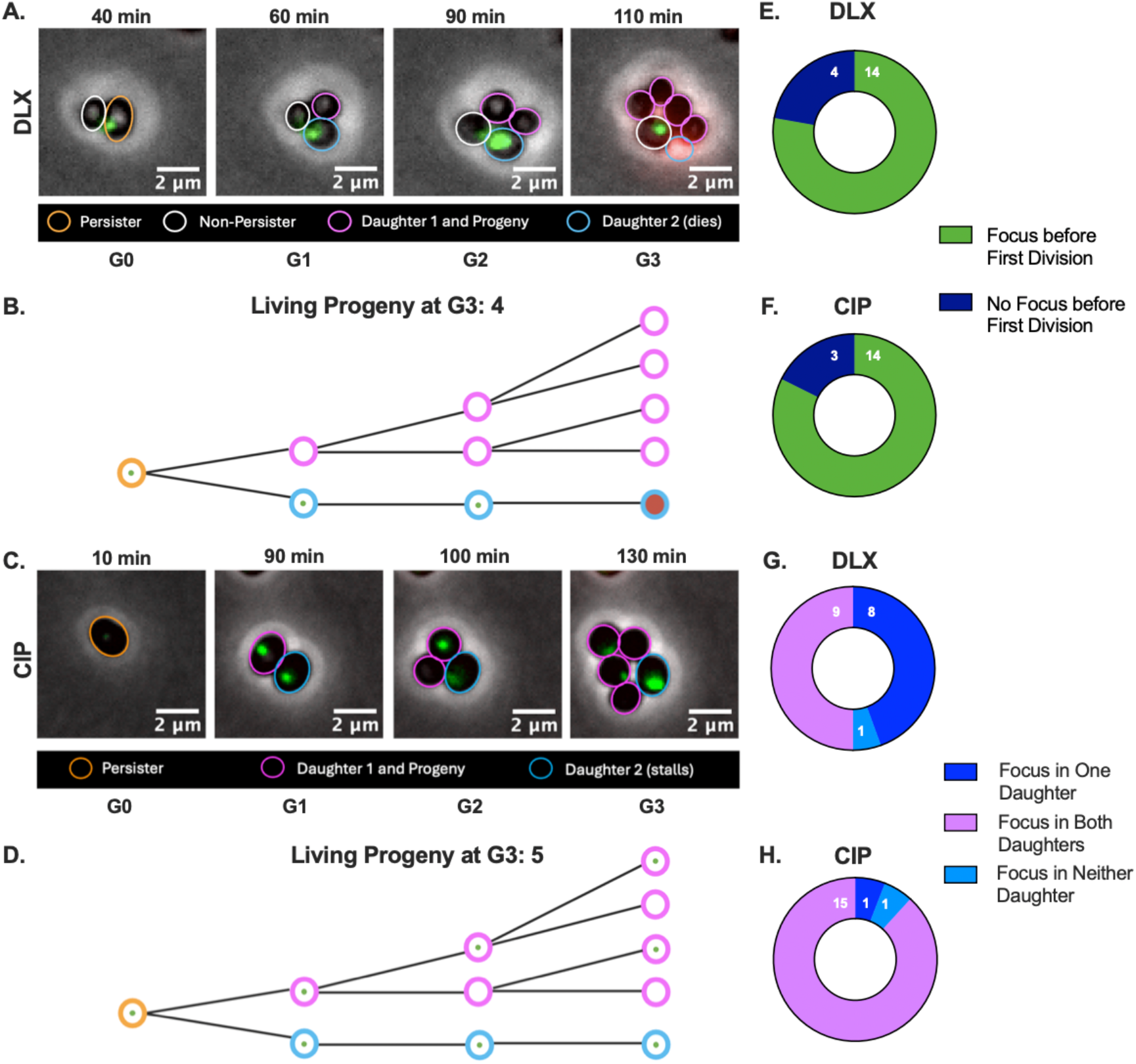
Both FQ-treated persisters and their progeny experience and respond to DSBs. (A-D) Stationary-phase HG003::*P*_*recA*_*-recA-gfp* was treated for 5 h with DLX or CIP then observed via timelapse microscopy. (A) A DLX non-persister (white outline) forms a focus and never divides. A DLX persister (orange outline, G0) forms a RecA-GFP focus then divides into two cells (blue and magenta, G1). Daughter 1 (magenta outline) does not inherit a focus and instead divides into two cells at G2, all of which divide again in G3; therefore, it appears that this daughter escaped FQ-induced damage. Daughter 2 (blue outline) inherits the focus, stalls, and lyses, suggesting that it inherited a DSB from the persister and failed to repair this damage. (B, D) Schematics illustrating the division events observable in (A) and (C), respectively, as well as the number of living progeny (defined as cells that had not taken up PI or lysed) at the end of G3. Circles represent cells, green dots represent RecA-GFP foci, and red shading represents PI staining. Created with BioRender. (C) A CIP persister (orange outline, G0) forms a faint RecA-GFP focus then divides into two cells (blue and magenta, G1), each of which has a focus. Daughter 1 (magenta outline) divides and passes its focus on to one G2 progeny. Both G2 progeny divide and produce one daughter with a faint focus. Daughter 2 (blue outline) never divides. (E-F) The proportion of all trackable (E) DLX and (F) CIP persisters that formed a focus before their first division was calculated. Compare to Fig. S7F. (G-H) The proportion of persisters that produced zero, one, or two daughters with a focus was calculated. Images are representative of at least three independent experiments.

Beyond persisters and their immediate progeny, we also often observed symptoms of DNA damage including RecA-GFP focus formation, swelling, stalling, and death, in later generations of the cohort **(Fig. S10;** compare to Fig. S7B-E) (49). These observations suggest that even the granddaughters of persisters experience some consequences of FQ treatment. We hypothesize that the inheritance of FQ molecules in the cytoplasm could cause topoisomerase trapping and subsequent DSB formation in later generations. While testing this hypothesis is beyond the scope of this work, we will explore this possibility in the future. In all, our data challenge the idea that persisters repair damage then produce normal, healthy progeny.

### Starvation after FQ treatment increases persistence

After finding that *S. aureus* persisters and their progeny engage RecA as they attempt to repair DSBs following FQ treatment, we sought to elucidate how nutrient availability during recovery impacts the success of persister resuscitation. To determine whether post-treatment nutrient limitation enhances FQ persistence in *S. aureus*, we treated WT *S. aureus* for 5 h with DLX or CIP then either plated the cells onto nutrient-rich media (Mueller-Hinton Agar [MHA]) immediately after antibiotic was removed or recovered the cells for 2 or 4 h in nutrient-free PBS to starve them before plating onto MHA.

We confirmed that starvation does not impact the culturability of untreated WT, *recA*::Tn, or *rexB*::Tn cells **(Fig. S11)**, and we found that starving WT populations for 2 h following DLX or CIP treatment significantly increased survival ∼2-3 fold. Extending the time of starvation to 4 h further increased survival to ∼10-fold **(Fig. 3A-B)**. To confirm that these results were not an artefact of our CFU assays, we also used timelapse microscopy to determine the fraction of cells that persisted in FQ-treated populations that were starved or immediately fed after treatment. These data demonstrate that the persister fraction in populations starved for 2 h following treatment was indeed significantly higher than in the fed populations **(Fig. S12; Movies S17-20)**.

**Figure 3.**
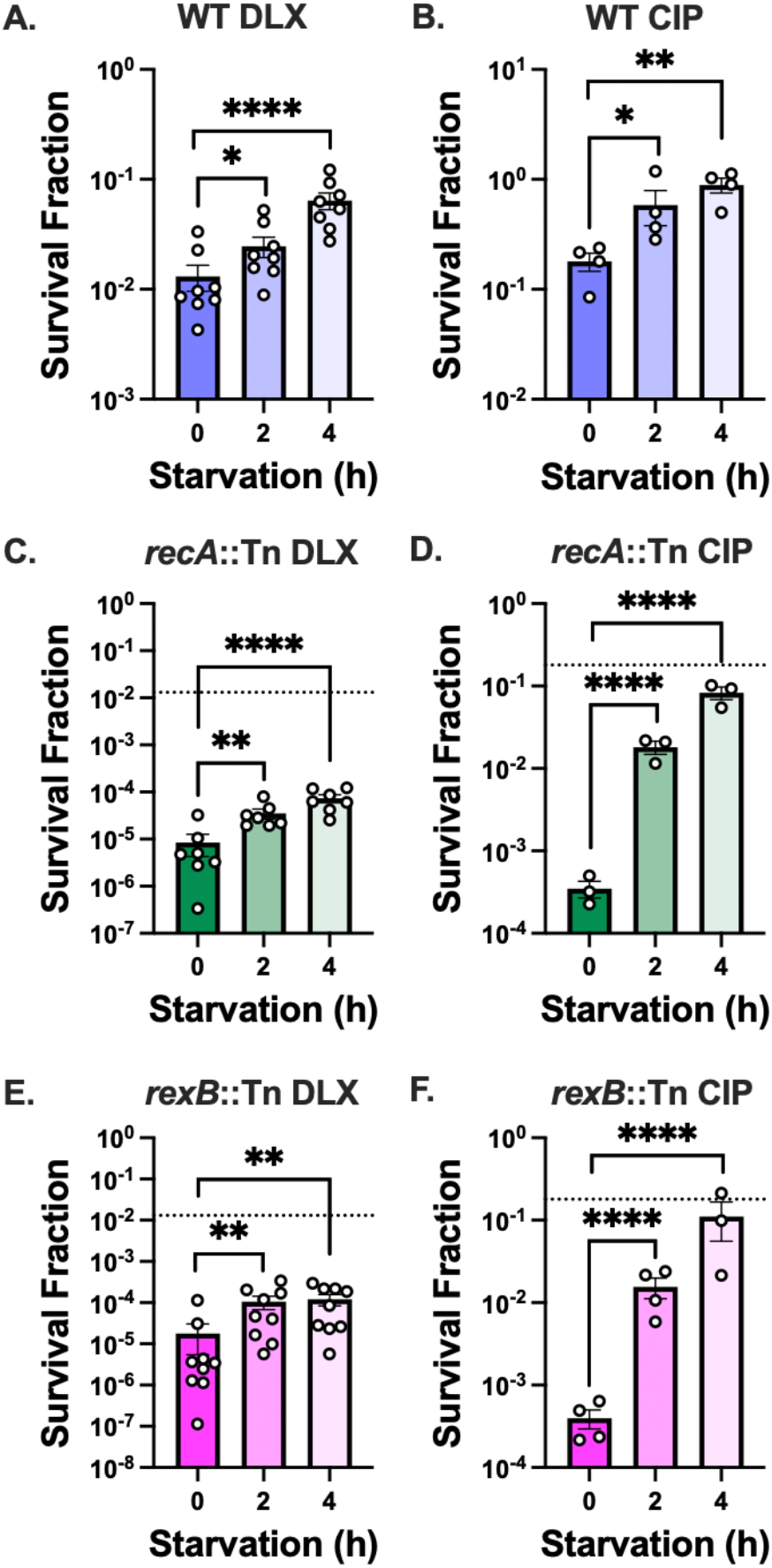
Starvation after DLX and CIP treatment increases persistence even in mutants lacking DSB-repair enzymes. Stationary-phase *S. aureus* (A-B) WT, (C-D) *recA*::Tn, and (E-F) *rexB*::Tn were treated with 5 μg/mL DLX or 10 μg/mL CIP for 5 h. After treatment ended, the cells were starved for 0, 2, or 4 h before being plated onto MHA for CFU enumeration. Dotted lines indicate the mean survival of non-starved WT cells treated with a given FQ. At least three independent replicates were performed for each experiment. *P* values were calculated using Dunnett’s multiple comparisons test following ANOVA to compare log-transformed survival fractions at each timepoint to 0 h. \**P <* 0.05, \*\**P <* 0.01, \*\*\*\**P <* 0.001. Error bars denote SEM.

### DSB-repair mechanisms are not essential for starvation-induced survival increase

Next, we asked whether DSB repair is necessary for the observed survival benefit caused by post-treatment starvation, as it is in *E. coli* (38). Remarkably, *S. aureus* lacking *recA* still showed significantly increased survival when starved after treatment with DLX (∼9-fold) or CIP (∼250-fold) for 4 h **(Fig. 3C-D)**. While we note that the effect of loss of *recA* on DLX survival could not be fully overcome by post-treatment starvation, starving CIP-treated *recA*::Tn for 4 h increased its survival to levels comparable to WT cells. These data suggest that the survival increase caused by post-treatment starvation is not dependent on RecA.

Since RecA-mediated HR is the only known DSB repair pathway in *S. aureus*, we hypothesized that *recA*::Tn employs some alternate DSB repair pathway during starvation to increase its survival (50). Therefore, we assessed the survival of mutants lacking *rexB. rexB* encodes an exonuclease required to process DNA ends at DSBs before the breaks can be repaired via RecA-mediated HR (50). Although *S. aureus* lacks the enzymes needed to repair DSBs via non-homologous end-joining (NHEJ), Ha and Edwards proposed that RexAB and ligase LigA could enable DSB repair via the alternative-end joining (A-EJ) pathway in *S. aureus*, though this has not been experimentally demonstrated (50, 51). Since RexB is required for both RecA-mediated HR and presumably, A-EJ, loss of *rexB* should eliminate *S. aureus*’s ability to repair DSBs via both A-EJ and HR.

We confirmed that loss of RexB function significantly decreased FQ persistence in stationary-phase *S. aureus*, similar to populations lacking *recA* **(Fig. S13)**. Surprisingly, like *recA*::Tn, *rexB*::Tn mutants benefitted from nutrient deprivation during recovery from DLX or CIP treatment, as starving for 4 h increased survival following DLX treatment ∼7-fold and CIP treatment ∼275-fold **(Fig. 3E-F)**. While we cannot rule out the roles of enzymes that repair non-DSB DNA lesions, these data indicate that DSB repair is not essential for the increased DLX and CIP survival following starvation, suggesting that post-treatment starvation increases survival by other means.

### Post-treatment starvation limits DSBs in persisters

To further investigate how starvation post-FQ treatment affects DNA damage and repair in *S. aureus* persisters, we monitored our RecA-GFP translational reporter strain as it recovered on Mueller-Hinton agarose following post-FQ treatment starvation **(Fig. 4A-D; Movies S17-20;** compare to **Fig. 2A-D** and **Movies S13-16)**. We found that starved cells were less likely to form foci before dividing. Indeed, 29% of starved DLX persisters divided without forming RecA-GFP foci before division **(Fig. 4G)**, compared to 22% of unstarved DLX persisters **(Fig. 2E)**. Similarly, 44% of starved CIP persisters divided before forming a focus **(Fig. 4H)**, compared to 18% of persisters from cultures that were fed right after treatment **(Fig. 2F)**. Furthermore, although 65% of starved DLX persisters still produced two daughters with foci **(Fig. 4I**; compare to **Fig. 2G)**, starvation decreased the percentage of CIP persisters that produced two daughters with foci from 88% to 50% **(Fig. 4J**; compare to **Fig. 2H)**. These data suggest that starvation makes it more likely that CIP persisters will produce at least one daughter that escapes treatment without developing a DSB, though this may not be true for DLX persisters.

**Figure 4.**
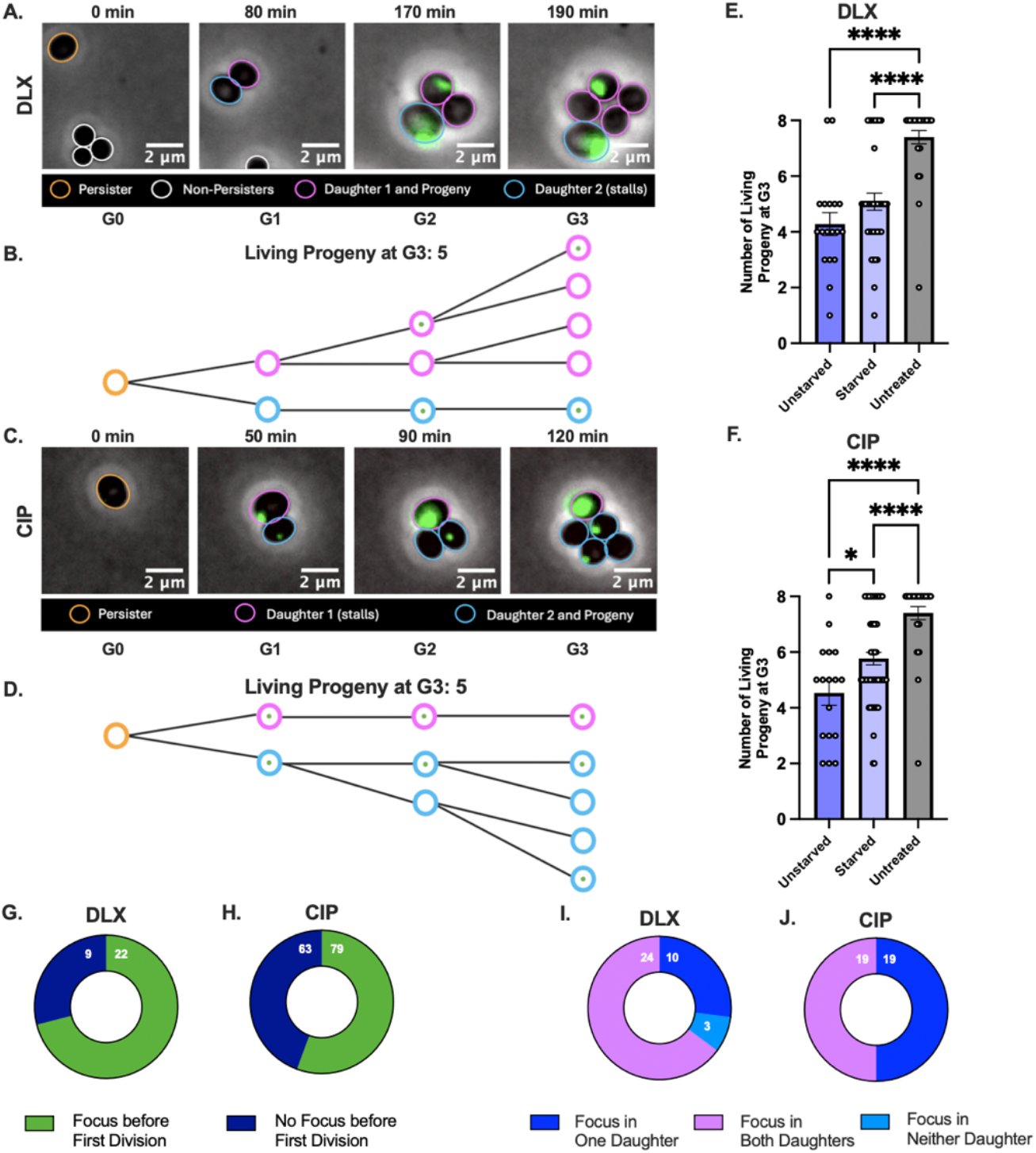
Post-treatment starvation reduces the appearance of DSBs in persisters and improves progeny success. (A-D) *S. aureus* HG003::*P*_*recA*_*-recA-gfp* was treated for 5 h with (A-B) DLX or (C-D) CIP, starved for 2 h, then tracked via timelapse microscopy in three independent imaging experiments. (A) A starved DLX persister (orange outline, G0) does not form a RecA-GFP focus before dividing into two cells at G1. Daughter 1 (magenta outline) divides into two cells, one of which forms a focus at G2. Both of these cells divide and produce two progeny, resulting in four magenta cells at G3. Daughter 2 (blue outline) forms a bright focus over time and stalls. (B, D) Schematics illustrating the division events observable in (A) and (C), respectively, as well as the number of living progeny (defined as not having taken up PI or lysed) at the end of G3. Circles represent cells, and green dots represent RecA-GFP foci. Created with BioRender. (C) A starved CIP persister (orange outline, G0) does not form a RecA-GFP focus before dividing into two cells at G1, both of which have a focus. The presence of a focus in both daughters indicates that both daughters are experiencing a DSB. The focus of daughter 1 (magenta outline) grows brighter, and it fails to divide. Daughter 2 (blue outline) divides, passing its focus on to one of its daughters in G2. Both blue daughters produce two more cells at G3, and one of each has a focus. (E-F) The number of living progeny at G3 for 30 untreated cells was compared with (E) 18 unstarved DLX persisters and 37 starved DLX persisters or (F) 17 unstarved CIP persisters and 56 starved CIP persisters. (G-H) The proportion of (G) all trackable starved DLX and (H) 143 randomly selected starved CIP persisters that formed a focus at any point before their first division was calculated. Compare to Fig. 2E-F. (I-J) The proportion of starved persisters that produced zero, one, or two daughters with a focus was calculated. Images are representative of at least three independent experiments.

To assess how post-treatment starvation impacts the success of persister progeny in later generations, we counted the number of living progeny at G3 for untreated cells, unstarved FQ persisters, and starved FQ persisters **(Fig. 4E-F)**. If all the progeny survived, there would be eight G3 daughter cells. We found that 73% of untreated cells produced the maximum eight cells at G3. Unstarved persisters usually produced only 4-6 living progeny by G3 due to death or stalling of a progeny at G1 or G2. Indeed, only 11% of unstarved DLX persisters and 6% of unstarved CIP persisters produced eight cells at G3. Following starvation, the number of DLX and CIP persisters giving rise to eight living progenies at G3 increased to 22% and 25%, respectively **(Fig. 4E-F; Fig. S14)**. Our data suggest that post-treatment starvation limits FQ-induced damage in both persisters and their progeny. Importantly, we see that starvation that extends into the post-FQ treatment period not only increases the number of persisters in each FQ-treated population but also augments the number of viable persister progenies in each persister-derived microcolony.

### Reduction of oxidative stress does not fully explain how starvation increases survival

After finding that post-treatment starvation increases survival by limiting damage rather than enhancing repair, we explored the mechanism whereby starvation limits damage. Our previous work suggests that nutrient stimulation during FQ treatment of stationary-phase *S. aureus* increases reactive oxygen species (ROS) generation, leading to enhanced cell death (20). Additionally, Hong and colleagues showed that ROS generation after treating *E. coli* with the quinolone nalidixic acid leads to cell death, and sequestering ROS after treatment increases *E. coli* survival (52). Therefore, we asked whether starvation allows FQ-treated *S. aureus* to cope with oxidative stress before resuming growth, thereby limiting damage and bolstering survival. If this model is applicable to *S. aureus*, we would expect that decreasing oxidative stress as cells recover from FQ treatment, even in the presence of nutrients, would increase survival.

To test this hypothesis, we treated *S. aureus* for 5 h with FQs then recovered the cells on nutritive agar under four conditions designed to limit oxidative stress: aerobically in the presence of **(1)** the antioxidant glutathione (GSH), **(2)** the iron chelator 2-2′-bipyridine (Bipy), **(3)** the antioxidant N-acetylcysteine (NAC), or **(4)** in an anaerobic chamber (53–56). Neither GSH nor Bipy appreciably increased the survival of WT, *recA*::Tn, or *rexB*::Tn **(Fig. S15A-D)**. For reasons unknown, NAC inhibited the growth of *recA*::Tn and *rexB*::Tn, so we only tested its effect on the recovery of WT and found significantly increased survival following CIP, but not DLX, treatment **(Fig. S15E-F)**. Recovering cells under anaerobic conditions failed to rescue any of the strains from DLX but increased the survival of all three strains treated with CIP **(Fig. S15G-H)**. However, since NAC and anaerobic nutritive recovery were not as effective as post-CIP treatment starvation at increasing survival, limiting oxidative stress is not sufficient to fully account for starvation-induced survival increase.

To further explore the role of oxidative stress, we generated strains incapable of producing one of three major ROS detoxification enzymes: catalase (*katA*::Tn), 2-cysteine peroxiredoxin (*tpx*::Tn), or superoxide dismutase (*sodA*::Tn) (57). If these genes contribute to starvation-induced survival increase, we would expect that disrupting them would preclude survival increase. We observed that starvation of *katA*::Tn and *tpx*::Tn led to survival comparable to WT **(Fig. S16A-D)**. However, while *sodA*::Tn had higher baseline survival than WT, its survival post-FQ treatment did not increase with starvation **(Fig. S16E-F)**. Therefore, SodA may contribute to starvation-induced survival increase, and the mechanism underlying its high baseline FQ survival will be explored in future work. Overall, our data suggest that starvation could increase CIP survival by decreasing oxidative stress, but the lack of effect on DLX-treated cells prompted us to seek alternative explanations.

### Post-treatment starvation may limit topoisomerase trapping during recovery

Next, we hypothesized that starvation may allow the cells to enzymatically free topoisomerases that were trapped during treatment before DSBs culminate. If this hypothesis is correct, we would expect that disrupting the function of XseAB, a homolog of ExoVII, which is known to free FQ-trapped topoisomerases in *E. coli*, would prevent starvation-induced survival increase (34). However, while our *xseA*::Tn mutant had lower DLX and CIP survival than WT, it still benefitted from post-treatment starvation **(Fig. S17)**. Therefore, we conclude that post-treatment starvation does not likely increase survival through enzymatic freeing of trapped topoisomerases.

We then considered whether post-treatment starvation may limit topoisomerase trapping during the post-treatment period. In stationary phase, *S. aureus* performs essentially no DNA synthesis (20). While stationary-phase cells maintain some transcriptional activity, it is much lower than under nutrient-replete conditions (20). Therefore, it is possible that most topoisomerase trapping occurs during post-treatment once cells encounter nutrients and topoisomerase activity increases, rather than during treatment itself. We hypothesized that post-treatment starvation allows the cells to expel FQs that have accumulated within the cell during treatment before nucleic acid synthesis and requisite topoisomerase activity resume, limiting topoisomerase trapping and subsequent DSBs.

To determine whether post-treatment starvation allows FQ expulsion, we used liquid chromatography-tandem mass spectrometry (LC-MS/MS) to measure intracellular abundance of DLX and CIP at the end of treatment, immediately after washing away extracellular FQ, and at 2 and 4 h of starvation (27). We found that the intracellular abundance of both DLX and CIP relative to a cloxacillin (CLX) spike-in control decreased significantly after the washing procedure **(Fig. S18-19)**. DLX abundance only decreased ∼2-3-fold throughout the starvation period **(Fig. S19A-C)**. These data are consistent with DLX’s reported ability to hyperaccumulate within *S. aureus* cells (58). By comparison, CIP abundance decreased ∼25-fold in WT and HR mutant strains, which could explain why we detected >200-fold increase in the survival of *recA* and *rexB* mutants following starvation post-CIP treatment **(Fig. S19D-F)**.

### Inhibiting nucleic acid synthesis after FQ treatment increases survival even when nutrients are present

After finding that post-treatment starvation may promote FQ expulsion, we sought to determine whether post-treatment starvation inhibits nucleic acid synthesis. We used ^3^H-uridine to label newly synthesized RNA and DNA in *S. aureus* during incubation in PBS (starved condition) or cation-adjusted Mueller-Hinton broth (CA-MHB; non-starved condition) following DLX or CIP treatment (20, 53, 59). We confirmed that starving cells for 2 h after FQ treatment decreased DNA synthesis in DLX-treated WT and *recA*::Tn and in all CIP-treated strains **(Fig. S20A-B)**. We also showed that 2 h of nutrient limitation decreased RNA synthesis, though not always significantly, in all strains following FQ treatment **(Fig. S20C-D)**.

To determine whether decreased nucleic acid synthesis was responsible for the increased survival of starved cells, we used chemical inhibitors to halt DNA and/or RNA synthesis after FQ treatment. We recovered DLX- and CIP-treated cells in CA-MHB containing either the transcription inhibitor rifampicin (RIF), the DNA synthesis inhibitor 5-fluoro-2’-deoxycytidine (FDC), or both RIF and FDC at doses we confirmed inhibited RNA or DNA synthesis, respectively, without killing the cells **(Fig. S21)**.

Inhibiting replication or transcription alone for 2 h was insufficient to rescue DLX-treated cells **(Fig. 5A-C)**. Instead, WT and *recA*::Tn could only be rescued from DLX by inhibiting both replication and transcription. Inhibiting both DNA and RNA synthesis failed to rescue *rexB*::Tn. Previous studies show that DLX targets both gyrase (relieves strain in the DNA during RNA and DNA synthesis) and topoisomerase IV (decatenates newly replicated chromosomes) to equal extents (23–26, 60, 61). We confirmed this finding by showing that strains with mutations in the quinolone resistance-determining regions of gyrase (GyrA S84L) or topoisomerase IV (ParC S80Y) exhibit equal increases in DLX MIC **(Fig. S22A)** (62). Therefore, it is not surprising that both DNA and RNA synthesis must be inhibited to rescue *S. aureus* from DLX treatment since inhibiting both these processes should decrease the demand for both topoisomerases targeted by DLX.

**Figure 5.**
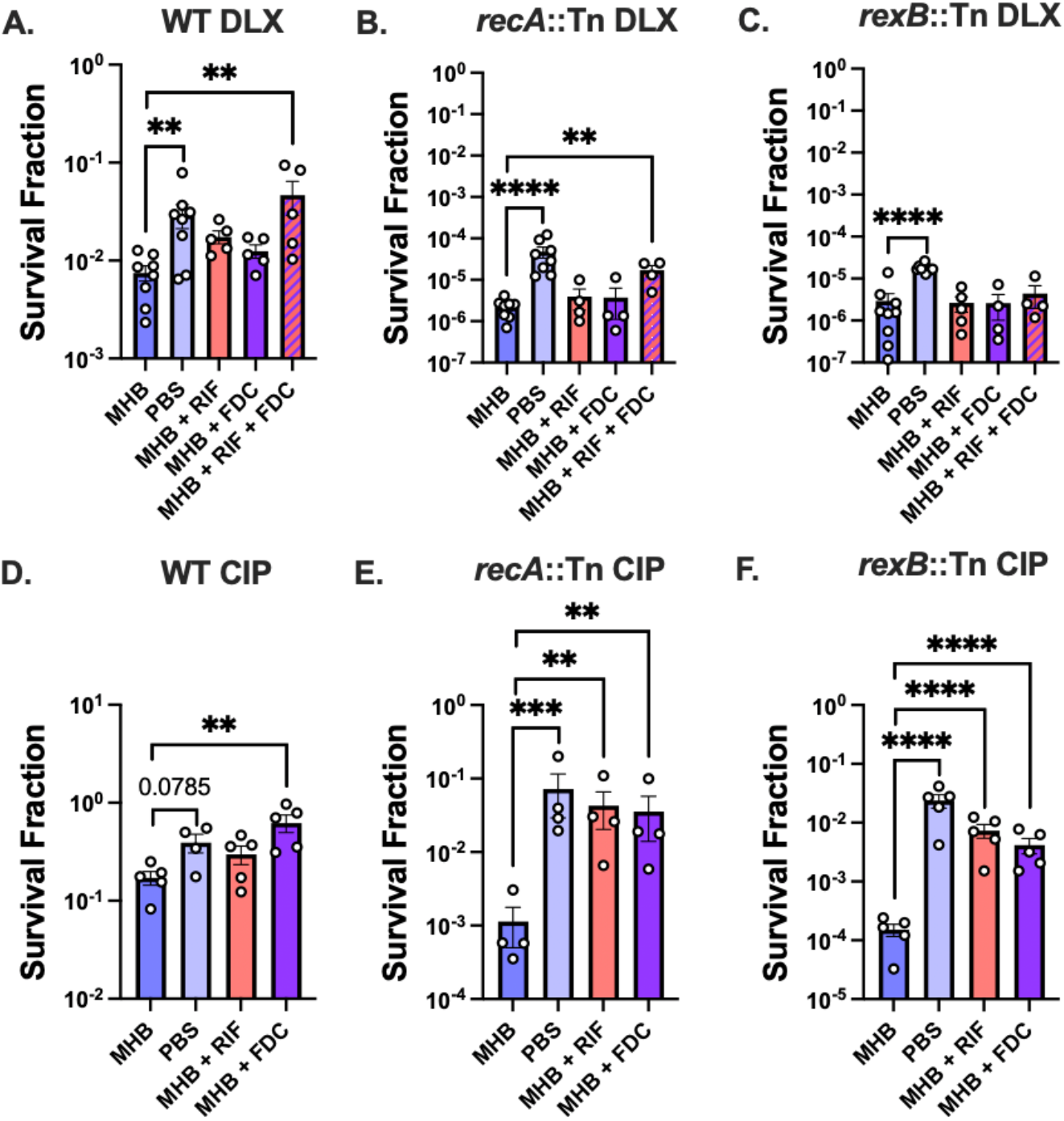
Inhibiting nucleic acid synthesis during nutritive recovery accounts for increased survival caused by post-treatment starvation. Stationary-phase *S. aureus* WT, *recA*::Tn, and *rexB*::Tn were treated with (A-C) 5 μg/mL DLX or (D-F) 10 μg/mL CIP. Then, cells were recovered for 2 h in MHB (not starved), PBS (starved), MHB + 0.02 μg/mL RIF, MHB + 1 μg/mL FDC, or MHB + 0.02 μg/mL RIF + 1 μg/mL FDC before plating onto inhibitor-free nutritive media to determine survival. *P* values were calculated using Dunnett’s multiple comparisons test following ANOVA to compare log-transformed survival fractions of each condition to the non-starved condition. \*\**P <* 0.01, \*\*\**P <* 0.005, \*\*\*\**P <* 0.001. At least four independent replicates were performed. Error bars denote SEM.

By comparison, we confirmed with our GyrA S84L and ParC S80Y mutants that CIP predominantly inhibits topoisomerase IV (**Fig. S22B**) (63–66). Consistent with this, we found that inhibiting DNA synthesis alone, which would circumvent the need for topoisomerase IV to decatenate sister chromosomes, improved the survival of CIP-treated WT. Inhibiting either DNA or RNA synthesis was sufficient to rescue CIP-treated *recA*::Tn and *rexB*::Tn **(Fig. 5D-F)**. In all, these results suggest that post-treatment starvation increases survival mainly by delaying the resumption of nucleic acid synthesis and decreasing activity of the topoisomerases targeted by the drugs.

## Discussion

Understanding the responses of bacterial persisters to environmental triggers and antibiotics during and after treatment is crucial for developing effective therapeutic strategies. In this study, we show that *S. aureus* persisters from stationary-phase cultures, which are likely representative of bacteria dwelling in nutrient-limited host sites, do not die or show evidence of DNA damage during FQ treatment (2–7). Instead, although FQs accumulate in cells throughout treatment, DSBs appear to culminate after treatment ends and nutrients are restored. This is reminiscent of recent studies showing that non-growing populations of FQ-treated *E. coli* and *S. enterica* remain viable in the presence of the drugs (39, 40). We note that this phenomenon is not generalizable to all bacterial species, as our previous study showed that starved *Pseudomonas aeruginosa* non-persisters die during levofloxacin treatment (53). For bacterial species that remain alive during antibiotic treatment, including our *S. aureus* populations, post-FQ treatment nutrient conditions and molecular events, such as damage repair processes and growth resumption, can impact death and persistence.

Given the significance of post-FQ treatment molecular events, identifying the enzymes and pathways that are important for persister recovery and resuscitation is crucial. In *E. coli* FQ persisters, enzymes involved in DNA break repair, including RecA, are vital for survival (33, 35, 67, 68). Here, we show that RecA- and RexB-mediated DSB repair is important for stationary-phase *S. aureus* FQ persistence, especially during the post-treatment recovery period, as previously shown for FQ-treated *S. aureus* in rich media (34). Similar to *E. coli*, we show that *S. aureus* persisters fed immediately after treatment do not usually survive by simply escaping FQ-induced DNA damage (35, 39). However, while *E. coli* persisters appear to repair damage before producing progeny, *S. aureus* persisters often pass on DNA damage to one or both daughter cells, reminiscent of what we recently observed in *P. aeruginosa* (69). Moreover, descendants of the original *S. aureus* persister that emerge after two to three generations frequently display evidence of FQ-induced damage. These data suggest that both *S. aureus* persisters and their progeny suffer some consequences from FQ treatment, which defies the paradigm of *E. coli* persisters from starved cultures that appear to complete DNA repair before cell division resumes, subsequently producing healthy daughter cells (39).

Previously, it was demonstrated that in *E. coli* and *S. enterica*, starving stationary-phase cells after FQ treatment increases survival (38, 40). Since *E. coli* populations lacking *recA* cannot be rescued by post-treatment starvation, it is thought that post-treatment starvation aids *E. coli* by allowing cells to perform RecA-mediated DSB repair before nutrient replenishment stimulates new DNA/RNA synthesis (38). Consistent with the *E. coli* model, we observed that starving DLX- and CIP-treated *S. aureus* after antibiotic removal enhanced persistence. However, in contrast to *E. coli*, post-treatment starvation increased *S. aureus* FQ persistence even in DSB-repair-deficient cells. In the case of CIP-treated *S. aureus* DSB repair mutants, starvation alarmingly increased survival >200-fold. These data suggest that DSB-repair mechanisms are not essential for *S. aureus* starvation-mediated survival increase. Instead, we observed that starvation allowed a large portion of persister cells to divide before forming RecA-GFP foci, suggesting that they divided before experiencing FQ-induced DSBs. Additionally, starved persisters were more likely to produce at least one daughter cell with no RecA-GFP focus, implying that this daughter inherited a chromosome that avoided FQ-trapped topoisomerases and DSBs. Our data suggest that *S. aureus* persisters may experience and manage DSBs from FQ treatment differently than the *E. coli* models. Additionally, we found that the production of reactive oxygen species and metabolites following FQ treatment may play a limited role in the demise of *S. aureus*, contrary to previous reports on *E. coli* (52). Our data emphasize the need to investigate persister survival strategies in different pathogens to deduce shared and unique responses in distinct species.

Based on our data, we present the following model illustrating how *S. aureus* persisters overcome FQs, which differs substantially from models based on other bacteria, namely *E. coli* **(Fig. 6)**. In stationary phase, most *S. aureus* cells have two completely replicated chromosomes (46). Our results show that FQs accumulate within the *S. aureus* cells during treatment. However, since nucleic acid synthesis is limited during stationary phase, there are fewer active topoisomerases for FQs to bind and trap during treatment. When nutrients are replenished immediately after treatment ends, many cells rapidly resume *de novo* DNA and RNA synthesis in response to the nutritional signals. This could allow FQs that accumulated intracellularly throughout treatment to trap active topoisomerases. Trapped topoisomerases can lead to DSBs that culminate either before division (in the parent persister cell) or after division (in one or both daughter cells). A daughter cell that inherits a chromosome with a trapped topoisomerase or DSB must repair this damage via RecA-, RexB-, or XseA-mediated processes to survive; failure to repair the damage leads to growth stalling and eventual death. Interestingly, we observed that most persisters from cells that were fed right after FQ treatment produce two progenies with DSBs, implying that both of their chromosomes had trapped topoisomerases.

**Figure 6.**
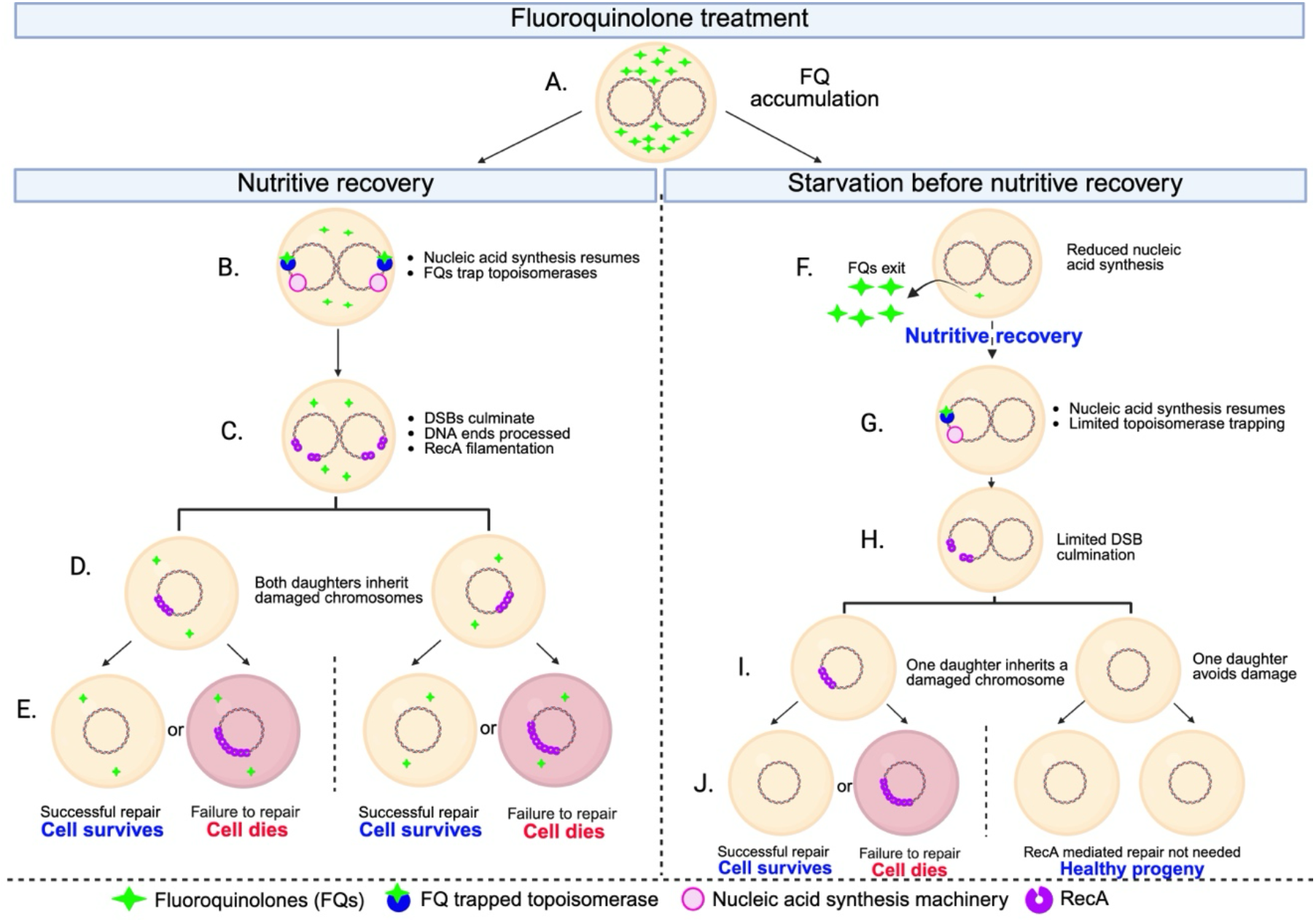
Schematic representation of events during S. aureus fluoroquinolone (FQ) treatment and recovery. **A.** During treatment, FQs accumulate within the cells. **B**. When nutrients are replenished immediately after treatment ends, cells rapidly resume nucleic acid synthesis, enabling accumulated FQs to trap topoisomerases. **C**. Topoisomerase trapping often leads to DSBs in both chromosomes. **D**. Consequently, both daughter cells inherit a chromosome with a culminated DSB. **E**. These daughters can either successfully engage RecA-mediated repair and survive or fail to repair the damage, resulting in stalling and eventual death. **F**. In contrast, starvation following treatment delays the resumption of nucleic acid synthesis. During this period, intracellular FQ levels decline. **G**. Once nutrients are restored after starvation, fewer residual FQ molecules remain to trap newly activated topoisomerases. **H**. As a result, fewer DSBs culminate. **I**. This appears to increase the likelihood that at least one daughter cell inherits an undamaged chromosome. **J**. This allows that daughter to survive independently of RecA. Created with BioRender.

With starvation following treatment, the resumption of nucleic acid synthesis is delayed, and the cells decrease the intracellular abundance of FQs—significantly for CIP but less for DLX. The decreased intracellular CIP concentration could explain how post-treatment starvation rescues survival almost fully even for *recA* and *rexB* mutants. While CIP is zwitterionic and can easily pass through the cell membrane, DLX is anionic at neutral cytoplasmic pH (58). DLX’s hyperaccumulating properties may explain its enhanced intracellular retention during starvation and likely prolong its bactericidal activity, even among starved cells. Based on our data, post-treatment starvation seems to allow more cells (particularly following CIP treatment) to pass on a chromosome with no trapped topoisomerase to at least one daughter, allowing it to survive without DSB repair proteins like RecA, since it does not experience a DSB. In all, we conclude that by delaying nucleic acid synthesis, post-treatment starvation increases the odds that an FQ-treated *S. aureus* cell can divide once before experiencing a DSB, making it more likely that at least one of its progenies will go on to produce a microcolony.

While our data offer insight into how *S. aureus* FQ persisters manage DSBs, some outstanding questions pertaining to features of persisters and cells that die remain. We do not yet know whether topoisomerase trapping occurs during FQ treatment or recovery, nor do we know whether the number and locations of irreversibly trapped topoisomerases differentiate cells that persist from those that succumb to treatment. Furthermore, the mechanism by which FQs are expelled under starvation conditions following FQ removal and whether heterogeneity in FQ abundance among individual cells post-treatment impacts survival/persistence remain to be deduced.

Our data show that starvation before, during, and after FQ treatment not only increases the number of persisters in the population but also increases the number of viable progenies that arise from each persister. This is particularly concerning for the treatment of *S. aureus* infections. To improve treatment outcomes, we should consider the development of therapeutics and other strategies to target critical DNA repair pathways that *S. aureus* persisters depend on to survive FQ treatment. Additionally, physical methods to improve circulation at infection sites could potentially stimulate *S. aureus* metabolism, precluding starvation-induced survival increase.

Our results offer a glimmer of hope, as they consistently show that it is more difficult for starved *S. aureus* to overcome the effects of the new FQ DLX than CIP at the tested doses. While post-treatment starvation led to nearly complete rescue of CIP-treated cells lacking key DSB-repair enzymes, the effect was much smaller for DLX-treated cells. The development of improved, dual-gyrase and topoisomerase IV-targeting FQs with better intracellular accumulation would enable us to combat recalcitrant *S. aureus* infections, even in nutrient-limited niches. Overall, our work identifies key genes and processes contributing to persistence in *S. aureus*, which could inform future clinically relevant strategies to combat this pathogen. Finally, further investigation into the specific nutrient requirements for *S. aureus* cells to succumb to FQ treatment could enable precision approaches to predict and enhance treatment success in patients.

## Materials and Methods

Experimental details on the following assays are included in *SI* Materials and Methods: growth curves, MIC determination, recovery under anaerobic and antioxidant conditions, quantification and inhibition of nucleic acid synthesis, and quantification of intracellular FQ abundance.

### Bacterial strains, plasmids, and primers

*S. aureus* strains used for all experiments in this study were derived from *S. aureus* HG003 **(Table S1)**. Plasmids and DNA primers used for strain construction and verification are listed in **Tables S2** and **S3**, respectively. Methods used for strain construction and verification are detailed in *SI* Materials and Methods.

### Survival assays

After 18 h of incubation, stationary-phase bacteria were treated with indicated doses of antibiotics. At designated times, 100 μL of each culture was collected, and cells were washed four times with PBS to dilute antibiotics to sub-MIC concentrations. Unless otherwise specified, cells were immediately serially diluted in PBS then plated onto Mueller-Hinton agar (MHA). Plates were incubated for 16-20 h before colonies were counted to determine CFU/mL after indicated lengths of treatment. Further details are provided in *SI* Materials and Methods.

### Starvation during recovery from antibiotics

*S. aureus* HG003, *recA*::Tn, *recA*::Tn [C], *rexB*::Tn, *rexB*::Tn [C], and *xseA*::Tn were grown for 18 h in 25 mL CA-MHB in 250-mL baffled flasks. After 18 h, cells were collected for plating and CFU enumeration. 2-mL cultures were treated with 5 μg/mL DLX or 10 μg/mL CIP for 5 h. After 5 h of FQ treatment, 1 mL of the treated cultures was collected and pelleted by centrifugation. Cells were washed four times with PBS before serial dilution and plating onto MHA. The remaining cells were left in PBS and incubated at 37 °C and 250 rpm for starvation during drug-free recovery. After 2 and 4 h, 10 μL of each sample was removed, serially diluted, and spotted onto MHA plates for CFU enumeration.

### Timelapse microscopy

*S. aureus* HG003::pCM29-P_*recA*_-*gfp* was grown for 18 h in CA-MHB containing 10 μg/mL chloramphenicol (CAM) then treated with indicated doses of DLX or CIP. HG003::*P*_*recA*_*-recA-gfp* was grown and treated similarly, except that CAM was not included. Following PBS washes, cells were either immediately imaged or starved for 2 h in PBS before imaging. Timelapse microscopy of starved and unstarved cells recovering from FQ treatment on MHA pads in a Bioptechs chamber was performed using a Zeiss Axiovert 200M microscope as described previously (62). Further details are available in *SI* Materials and Methods.

### Statistics

Statistical analyses were performed using GraphPad Prism 10.1.0 and Microsoft Excel 16.107.3. For all survival experiments, log_10_-transformed survival fractions were compared. When only two groups were being compared, F-tests were used to determine whether variance was equal between the two groups. When F < 0.05, we concluded that the groups had unequal variances. Then, two-tailed *t-*tests assuming equal or unequal variances (depending on the results of the F-tests) were used to calculate *P* values.

For comparisons of three or more groups, analysis of variance (ANOVA) was used to detect differences. Then, Dunnett’s multiple comparisons test was used to compare each condition to every other condition or to a control, as described in the figure captions. Since the MIC data in Fig. S22 showed very little variance, we used the nonparametric Kruskal-Wallis test followed by Dunn’s multiple comparisons test in this case. *P* < 0.05 was considered statistically significant throughout this study, and at least three independent biological replicates were performed for each experiment unless stated otherwise.

## Acknowledgments

The following reagents were provided by the Network on Antimicrobial Resistance in *Staphylococcus aureus* (NARSA) for distribution by BEI Resources, NIAID, NIH: *Staphylococcus aureus* subsp. *aureus*, Strain JE2, NR-46543; Transposon Mutant NE805 (SAUSA300_1178), NR-47348; Transposon Mutant NE1012 (SAUSA300_0869), NR-47555; Transposon mutant NE458 (SAUSA300_1472, NR-47001; Transposon Mutant NE1332 (SAUSA300_1659), NR-47874; Transposon Mutant NE1336 (SAUSA300_1232), NR-47908; Transposon Mutant NE1932 (SAUSA300_1513), NR-48474. We thank Dr. Brian Conlon for providing *S. aureus* HG003, Dr. Richard Novick for *S. aureus* RN4220 and phage Φ11, Dr. Francis Alonzo III for *S. aureus* RN9011 and pJC1111, Dr. Paul Fey for pBK123, and Dr. Alexander Horswill for pCM29.

We thank the following individuals at UConn Health for their invaluable assistance in conducting experiments: Kevin Higgins and Rob Speers (Office of Radiation Safety) and Dr. Yi Wu and Susan Staurovsky (Center for Cell Analysis and Modeling Microscopy Facility). We thank Dr. Jeremy Balsbaugh and Dr. Sonam Tamrakar (UConn Proteomics and Metabolomics Facility). We also thank the members of the Mok Lab—Dr. Patricia Hare, Travis LaGree, Angela Power, Juliet González, Jipeng Yue, Dr. Anna Truong, Dr. Ranjuna Weerasekera, and Nathaniel Rodney—for helpful discussions.

Our work was supported by funding awarded to W.W.K.M. from the University of Connecticut start-up fund and the National Institutes of Health (NIH; 1DP2GM146456-01, 4DP2AI138154). The funders had no role in the design of our experiments or the preparation of this manuscript.

