## Supporting Information Text and Figures for "Post-fluoroquinolone treatment molecular events and nutrient availability modulate *Staphylococcus aureus* antibiotic persistence"

**This PDF file includes:**

Supporting Information Text
*SI* Materials and Methods
Figures S1 to S22
Tables S1 to S4
Legends for Movies S1 to S20
*SI* References

**Other supporting materials for this manuscript include the following:**

Movies S1 to S20

### *SI* Materials and Methods

#### Bacterial strains, plasmids, and growth conditions.

*S. aureus* HG003 was used throughout this study as a wild-type strain and for genetic manipulations (1). Whole-genome sequencing was performed (SeqCenter) followed by single-nucleotide polymorphism (SNP) calling using Snippy to confirm the identity of this strain (2). Throughout this project, we used BioCyc and AureoWiki to obtain sequence information (3, 4). The strains *recA*::Tn (*recA* transposon mutant), *rexB*::Tn (*rexB* transposon mutant), and *xseA*::Tn (*xseA* transposon mutant) were created by using phage transduction to move transposon insertions from the respective JE2 transposon mutants from the Nebraska Transposon Mutant Library into HG003 following a method described earlier (5–7). *recA*::Tn [C] (*recA*::Tn with $P_{recA}$-*recA* complemented into the genome at the *attB* site) and *rexB*::Tn [C] (*rexB*::Tn with $P_{rexB}$-*rexB* complemented into the genome at the *attB* site) were constructed by integrating these constructs into a *S. aureus* pathogenicity island in HG003 using a previously described method (8, 9). *S. aureus* HG003::$P_{recA}$-*recA*-*gfp* (translational fusion of RecA and GFP used as a fluorescent reporter of DNA damage) was constructed similarly. Successful transposon insertion and complementation were confirmed by colony PCR (cPCR) using the primers listed in **Table S3**.

Bacterial strains were preserved in 25% glycerol and stored at -80 °C. For all experiments, bacteria were inoculated from glycerol stocks into 25 mL of cation-adjusted Mueller-Hinton broth (CA-MHB; BD Difco) in 250-mL baffled flasks and grown for 18 h at 37 °C and 250 rpm. For anaerobic experiments, reagents and materials such as pipettes, pipette tips, microcentrifuge tubes, PBS, and agar plates were incubated in a Coy Lab anaerobic chamber overnight before the experiment to deoxygenate the materials.

#### Chemicals.

Antibiotic stocks were prepared as follows: 10 mg/mL delafloxacin (DLX; MedChem Express) was dissolved in dimethyl sulfoxide (DMSO; Fisher Scientific), 10 mg/mL ciprofloxacin (CIP; Alfa Aesar) in 0.1 N HCl, and 20 mg/mL 5-fluoro-2'-deoxycytidine (FDC; Sigma Aldrich) in DMSO. These antibiotic stocks were stored at -20 °C and diluted to desired concentrations when added to bacterial cultures. Chloramphenicol (CAM; 10 mg/mL; Fisher Scientific) and erythromycin (ERM; 10 mg/mL; Acros Organics) were dissolved in molecular biology-grade absolute ethanol (Fisher Scientific) and stored at 4 °C. Anhydrotetracycline (100 µg/mL; Fisher) was dissolved in absolute ethanol and stored at -20 °C. Fresh 100 mg/mL stocks of ampicillin in water were made immediately before making selection media. Fresh 10 mg/mL stocks of rifampicin (RIF; TCI Chemicals) were made in DMSO before each experiment. Cadmium chloride ($CdCl_2$; 100 mM, Fisher Scientific) and sodium citrate (500 mg/mL or 0.02 M, depending on the procedure; Fisher) were dissolved in sterile water and stored at 4 °C. 2-2'-bipyridine (Bipy; 300 mM; Sigma Aldrich) was dissolved in absolute ethanol, glutathione (GSH; 75 mM; Fisher Scientific) was dissolved in water, and N-acetylcysteine (NAC; 100 mM; Thermo Scientific) was dissolved in ethanol; these were stored at room temperature until used. Phosphate-buffered saline (PBS) was prepared from PBS tablets (Fisher Scientific) and autoclaved before use. $^{3}$H-uridine (Revvity) was stored and handled according to the instructions from UConn Health's Office of Radiation Safety.

#### Making a cadmium-inducible *recA* expression system.

To place *recA* under the control of an inducible system, we cloned *recA* downstream of a cadmium-inducible promoter ($P_{cad}$) in pBK123 (10, 11). To construct *S. aureus* pBK123-$P_{cad}$-*recA*, *recA* was amplified from *S. aureus* HG003 using the primers *recA*-pBK123-F and *recA*-pBK123-R **(Table S3)**. Restriction sites for SalI and XmaI were added to the 5' ends of the primers. Then, the amplicon and pBK123 were digested for 2 h at 37 °C with the restriction enzymes SalI and XmaI (NEB) in rCutSmart buffer (NEB) using the manufacturer's protocol. Digested pBK123 and *recA* were ligated using T4 DNA ligase (ThermoFisher) following the manufacturer's method. The ligated products were transformed into chemically competent *E. coli* DH5α (Zymo Research), and transformants were confirmed by cPCR using primers pBK123_seq_F and *recA*_pBK123_R **(Table S3)** and Sanger sequencing (Azenta). $P_{cad}$-*recA* was electroporated into *S. aureus* RN4220 competent cells prepared as described previously, and transformants were selected by plating onto tryptic soy agar (TSA) containing 10 µg/mL CAM (5, 12–14). Transfer of the plasmid into *S. aureus* HG003 *recA*::Tn was performed using phage transduction as described previously (5–7).

**Making a *recA* transcriptional reporter.**

To construct a fluorescent transcriptional reporter of *recA,* the *recA* promoter (~500 bp immediately upstream of *recA*) was amplified from *S. aureus* HG003 genomic DNA. Restriction sites NheI and KpnI were added to the promoter using the primers pCM29_*recA*_F and pCM29_*recA*_R **(Table S3),** allowing it to be cloned into pCM29 at these sites (15). Clones were verified via whole-plasmid sequencing (Eurofins Genomics) before plasmids were transformed into *S. aureus* RN4220, then phage transduced into HG003.

**Making a *recA* translational reporter.**

To construct a chromosomal *recA* translational reporter, pCM29 containing *gfp* was linearized using the primers pCM29_Gibson_F and pCM29_Gibson_R **(Table S3)**. The *PrecA-recA* fragment was amplified from HG003 genomic DNA using the primers $P_{recA}$_Gibson_F and $P_{recA}$_Gibson_R. In addition to adding overlap sequences for Gibson assembly into pCM29 upstream of *gfp,* these primers added the sequence for a Gly-Gly-Ala-Ala-Gly linker to connect RecA with GFP **(Table S3)** (16). The linearized vector (pCM29-*gfp*) and the insert ($P_{recA}$-*recA* + linker + overlap) were assembled using Gibson assembly (NEB) according to the manufacturer's protocol and transformed into *E. coli* DH5α, followed by selection on LB agar plates containing 100 µg/mL ampicillin. Positive clones were confirmed by whole-plasmid sequencing. Subsequently, the $P_{recA}$-*recA-gfp* fragment was amplified from a confirmed pCM29 construct using the primers pJC1111_*recA*_Gibson_F and pJC1111_*gfp*_R **(Table S3)**. The pJC1111 vector was linearized by digestion with BamHI, and the amplified fragment was cloned into pJC1111 via Gibson assembly. The resulting constructs were verified by whole-plasmid sequencing. The confirmed pJC1111-$P_{recA}$-*recA-gfp* plasmid was then mini-prepped and electroporated into *S. aureus* RN9011. Finally, integration into *S. aureus* HG003 WT was achieved via phage transduction as described below.

**Verifying transposon mutants.**

The JE2 transposon mutant strains (*recA*::Tn, *rexB*::Tn, *xseA*::Tn, *katA*::Tn, *sodA*::Tn, and *tpx*::Tn) were obtained from the Nebraska Transposon Mutant Library through BEI Resources. We confirmed that these strains had transposon insertions in the correct locations via PCR as follows: *recA*::Tn using primers SA_*recA*_int_R and Buster, *rexB*::Tn using primers *rexB*_Tn_F and Upstream, *xseA*::Tn using primers *xseA*_Tn_F and Upstream, *sodA*::Tn using primers *sodA*_F and Upstream, *katA*::Tn using primers *katA*_F and Buster, *tpx*::Tn using primers *tpx*_F and Upstream **(Table S3)** (17). Transposon insertions were transduced into *S. aureus* HG003 to create HG003 *recA*::Tn, *rexB*::Tn, *xseA*::Tn, *sodA*::Tn, *katA*::Tn, and *tpx*::Tn mutants via phage transduction as described below.

**Constructing complementation strains.**

To complement $P_{recA}$-*recA* and $P_{rexB}$-*rexB* into HG003 *recA*::Tn and *rexB*::Tn, pJC1111 was digested with BamHI for 2 h at 37 °C. $P_{recA}$-*recA* was amplified from *S. aureus* HG003 genomic DNA using the primers pJC1111_*recA*_Gibson_F and pJC1111_*recA*_Gibson_R **(Table S3)**. $P_{recA}$-*recA* was then inserted into pJC1111 using Gibson assembly, and the product was transformed into *E. coli* DH5α competent cells. Transformants were confirmed by cPCR using the primers pJC1111_seq_F and pJC1111_seq_R before Sanger sequencing using the same primers **(Table S3)**. pJC1111-$P_{recA}$-*recA* was then electroporated into *S. aureus* RN9011 electrocompetent cells followed by selection on TSA containing 100 µM $CdCl_2$ + 10 µg/mL CAM. The genomic complementation of $P_{rexB}$-*rexB* into *rexB*::Tn was also performed using this method. $P_{rexB}$-*rexB* was amplified from the genomic DNA of *S. aureus* HG003 using the primers pJC1111_*rexB*_Gibson_F and pJC1111_*rexB*_Gibson_R **(Table S3).** Genomic complementation of *recA* and *rexB* into *S. aureus* HG003 *recA*::Tn and *rexB*::Tn were performed using phage transduction from RN9011 followed by selection on TSA plates containing 100 µM $CdCl_2$ + 500 µg/mL sodium citrate. The genomic complementation was confirmed by PCR using primers SaPI_chrom_verify and SaPI_vector_verify **(Table S3)** (18).

**Making topoisomerase mutants.**

To generate topoisomerase mutants, we amplified *gyrA* from *S. aureus* JE2's genomic DNA (which has a GyrA S84L mutation) using primers pJB38_*gyrA*_F and pJB38_*gyrA*_R that added an EcoRI restriction site upstream and a KpnI restriction site downstream of *gyrA*. We amplified *parC* with the S80Y mutation from *S. aureus* JE2's genomic DNA using primers pJB38_*parC*_F and pJB38_*parC*_R, flanking the gene with EcoR1 and KpnI restriction sites. The plasmid pJB38 and amplified *gyrA* and *parC* were then digested using EcoRI and KpnI restriction enzymes, ligated using quick ligase (NEB), and selected on LB agar containing ampicillin (19). The clones were confirmed by cPCR and whole-plasmid sequencing. Plasmids were electroporated into *S. aureus* RN4220, and transformants were selected on 10 µg/mL CAM at 30 °C to allow plasmid replication. Next, we propagated phage Φ11 on *S. aureus* RN4220 containing pJB38 with mutated *gyrA* or *parC* then transduced the mutations into *S. aureus* HG003. Clones were selected on TSA + 10 µg/mL CAM at 30 °C. *S. aureus* HG003 containing pJB38-*gyrA* or pJB38-*parC* was then streaked onto TSA + 10 µg/mL CAM and grown at 44 °C to force recombination. The clones were then passaged five times in TSB at 30 °C without any antibiotic to enable the second recombination and plasmid loss. After passaging the cells, the cultured clones were diluted up to $10^{-7}$ and plated on TSA with 100 ng/mL anhydrotetracycline to select for colonies that had lost the plasmid. The colonies from the plates were then replica patched on selective media. The potential *gyrA* mutants were grown on TSA, TSA + 0.5 µg/mL CIP, TSA + 10 µg/mL CIP and TSA + 10 µg/mL CAM. The potential *parC* mutants were grown on TSA, TSA + 2 µg/mL CIP, TSA + 25 µg/mL CIP and TSA + 10 µg/mL CAM (20). For potential *gyrA* mutants, we selected colonies that grew on TSA and TSA + 0.5 µg/mL CIP but did not grow on TSA + 10 µg/mL CIP or CAM. For potential *parC* mutants, we selected colonies that grew on TSA and TSA + 2 µg/mL CIP but did not grow on TSA + 25 µg/mL CIP or CAM. Finally, the DNA was extracted from these colonies for cPCR. *gyrA* was amplified using pJB38_*gyrA*_F and pJB38_*gyrA*_R, and *parC* was amplified using pJB38_*parC*_F and pJB38_*parC*_R. Amplicons were sent for Sanger sequencing using pJB38_*gyrA*_F + pJB38_*gyrA*_R (for *gyrA*), and pJB38_*parC*_F + *parC*_seq (for *parC*).

**Phage propagation and transduction.**

Desired gene constructs were transduced from *S. aureus* RN4220 or RN9011 into *S. aureus* HG003 WT, *recA*::Tn, or *rexB*::Tn using phage Φ11 following a method described previously (5–7). Φ11 propagated on *S. aureus* RN4220 or RN9011 containing the desired gene construct was harvested then used for transduction into our strains of interest. A loopful of heavily streaked overnight culture of the recipient strain was resuspended in TSB containing 5 mM $CaCl_2$ using a sterile loop. Then, 500 µL of resuspended bacterial culture was added to a 50-mL conical tube containing 1.5 mL TSB + 5 mM $CaCl_2$ and 500 µL harvested Φ11 (~$10^{10}$ plaque-forming units [PFU]/mL). The tubes were incubated at 37 °C for 20 min with shaking at 225 rpm. After 20 min, 1 mL ice-cold 0.02 M sodium citrate was added to the tube to halt phage binding by chelating $CaCl_2$. The tubes were then centrifuged for 10 min at 4,000 *x g* at 4 °C. The supernatant was removed, and the pellet was resuspended in 1 mL ice-cold 0.02 M sodium citrate. 100 µL of the resuspended culture was plated onto TSA plates containing 500 µg/mL sodium citrate and 10 µg/mL CAM, 10 µg/mL ERM, or 100 µM $CdCl_2$. The plates were incubated overnight at 37 °C. The following day, colonies were passaged by streaking onto new plates containing 500 µg/mL sodium citrate and the appropriate antibiotic for selection.

**Minimum inhibitory concentration (MIC) assays.**

The MICs of antibiotics were determined using the broth microdilution method (21). *S. aureus* strains were cultured overnight from glycerol stocks for 18 h before being diluted to $OD_{600}$ ~0.01 in fresh CA-MHB and grown to exponential phase ($OD_{600}$ ~0.2–0.4). Exponential-phase cultures were then diluted to $OD_{600}$ ~0.0001 (~$5x10^5$ cells/mL) in 5 mL CA-MHB. Two-fold serial dilutions of each antibiotic were prepared in 100 µL CA-MHB in a 96-well flat-bottom microtiter plate, and 100 µL of bacterial culture was added to each well. Antibiotic-free wells were included as negative controls. The plates were sealed with Breathe-Easy membranes (Millipore Sigma) and incubated for 18 h at 37 °C. After 18 h, $OD_{600}$ was measured using a BioTek Synergy plate reader, and the minimum antibiotic concentration that inhibited ~90% of bacterial growth was considered the MIC. The full range of MIC values obtained for each drug and each strain is reported in **Table S4**.

**Growth curves.**

*S. aureus* HG003 and *recA*::Tn were inoculated into 25 mL CA-MHB in 250-mL baffled flasks and grown overnight for 18 h at 37 °C and 250 rpm. After 18 h, 2 mL of overnight culture was dispensed into a test tube, and CFU/mL was tracked for an additional 24 h. 10 µL of the culture was removed at different time points, serially diluted in a 96-well microtiter plate containing 90 µL PBS, and then spotted onto MHA plates. The spots were allowed to dry before the plates were incubated overnight at 37 °C followed by colony-forming unit (CFU) enumeration. Survival fractions were calculated by dividing CFU/mL at the stated time points by CFU/mL at 0 h.

**FQ dose-dependent survival assays.**

*S. aureus* HG003 and HG003 *recA*::Tn were inoculated in 25 mL CA-MHB in 250-mL baffled flasks and grown overnight for 18 h at 37 °C and 250 rpm. After 18 h, 10 µL of each culture was collected for serial dilution and plating to enumerate CFU/mL. Then, 2-mL aliquots of the overnight cultures were dispensed into glass tubes and treated with varying concentrations of DLX (1, 5, and 10 µg/mL) or CIP (5, 10, 50, and 100 µg/mL) for 5 h at 37 °C, with shaking at 250 rpm. After 5 h, 1 mL of each culture was transferred to microcentrifuge tubes and centrifuged for 3 min at 21,000 *x g* before 900 µL of supernatant was removed. 900 µL sterile PBS was then added to the tube before the pellet was resuspended and pelleted again by centrifugation. This washing step was performed four times to dilute antibiotics to sub-inhibitory concentrations. 10 µL of each washed culture was then serially diluted in a 96-well round-bottom microtiter plate containing 90 µL PBS then spotted onto MHA plates. The spots were allowed to dry before the plates were incubated overnight in a 37 °C incubator. Survival fractions were calculated by dividing the CFU/mL at 5 h by the CFU/mL at 0 h.

**Time-dependent survival assays.**

*S. aureus* HG003 and HG003 *recA::*Tn were grown as described above. 2-mL cultures were then treated with 5 µg/mL DLX or 10 µg/mL CIP for 24 h. 100-µL samples were removed at indicated timepoints then centrifuged and washed as described above. The cultures were spotted onto MHA plates, and survival fractions were determined as described above.

**Inducible *recA* survival assay.**

*S. aureus* HG003 with pBK123, HG003 *recA*::Tn with pBK123, and *recA::*Tn with pBK123-$P_{cad}$-*recA* were grown for 18 h in CA-MHB containing 10 µg/mL CAM as described above. 10 µL of each culture was collected for serial dilution and CFU enumeration before antibiotic treatment. 2-mL aliquots of each culture were treated with 5 µg/mL DLX for 5 h then washed as described above. After washing, serially diluted cultures were spotted onto MHA plates containing 10 µg/mL CAM with or without 100 nM $CdCl_2$ for *recA* induction. The plates were incubated overnight at 37 °C before survival fractions were calculated.

**Survival assay with nucleic acid synthesis inhibitors.**

To ensure that RIF and FDC do not affect the survival of non-FQ treated *S. aureus*, 500 µL of stationary-phase cultures resuspended in fresh CA-MHB were dispensed into tubes containing 0.02 nmol/mL uracil and treated with either 0.02 µg/mL RIF, 1 µg/mL FDC, or no inhibitor for 2 h. At 0 and 2 h, 100 µL of each sample was removed, washed in PBS as described above, serially diluted, and spotted onto MHA. Colonies were counted following overnight incubation, and survival fractions were determined. Furthermore, we tested whether inhibiting nucleic acid synthesis after FQ treatment impacts *S. aureus* survival. *S. aureus* cultures were grown and treated with 5 µg/mL DLX or 10 µg/mL CIP for 5 h as described above. After treatment, the cells were washed and resuspended in PBS or CA-MHB. Cells that were resuspended in CA-MHB were treated with 0.02 µg/mL RIF, 1 µg/mL FDC, or both 0.02 µg/mL RIF and 1 µg/mL FDC for 2 h at 37 °C with shaking. After 0 and 2 h of incubation, the cells were pelleted by centrifugation (21,000 x g, 3 min) and washed to remove the nucleic acid synthesis inhibitors then serially diluted and spotted onto MHA for survival fraction determination.

**Recovery under anaerobic and antioxidant conditions.**

After treatment and washing as described above, *S. aureus* cells were spotted onto MHA with or without Bipy (300 µM), GSH (750 µM), or N-acetylcysteine (NAC; 5 mM and 10 mM) and incubated overnight (5, 22–24). MHA plates without antioxidants were supplemented with ethanol to account for the concentration of ethanol used to dissolve Bipy. For anaerobic recovery, the cells were recovered in an anaerobic chamber (Coy Laboratories; connected to 5% $CO_2$, 10% $H_2$, and balanced nitrogen-containing cylinders) on MHA that had been deoxygenated overnight. After overnight recovery, CFU/mL and survival fractions were calculated.

**Quantification of nucleic acid synthesis.**

We quantified nucleic acid synthesis to determine how starvation after antibiotic treatment affects nucleic acid synthesis using a previously described method with some modifications (25–27). Briefly, stationary-phase *S. aureus* was treated with FQs for 5 h as described above. Two aliquots were treated with each antibiotic. After 5 h, 1.7 mL of each culture was washed four times with PBS, and cells were resuspended in CA-MHB or PBS. Then, 1.5 mL of the resuspended cultures was added to conical tubes containing 1.5 µCi (1.5 µL of Revvity Stock) $^3$H-uridine and incubated at 37 °C and 250 rpm for 2 h.

To quantify total nucleic acid synthesis, 250 µL of the sample that had been incubated with $^3$H-uridine was added to a microcentrifuge tube containing 675 µL of 10% ice-cold trichloroacetic acid (TCA; Fisher Scientific) and incubated for at least 30 min on ice to precipitate total nucleic acids. To quantify DNA synthesis, another 250 µL of the sample was added to a microcentrifuge tube containing 27.5 µL of 3M KOH (Fisher Scientific) and incubated overnight at 37 °C without shaking to degrade alkali-labile RNA. After overnight incubation, ice-cold 10% TCA was added to the KOH-treated samples before incubation for at least 30 min on ice.

To remove unincorporated $^3$H-uridine and isolate radiolabeled nucleic acids, 25-mm Whatman GF/C binder-free microfiber glass filters (Cytiva) were soaked in 1 mL 10% TCA and overlaid on a Buchner funnel attached to a vacuum tube. The TCA-precipitated samples were applied onto the center of the filters and washed with 2 mL of ice-cold 10% TCA followed by 2 mL of ice-cold 70% ethanol. The filters were then allowed to dry for at least 2 h at room temperature.

After drying, the filters were transferred to 5 mL of Econo-Safe scintillation fluid (Fisher Scientific) in 6-mL scintillation vials (Fisher Scientific). A filter without a sample was added to one vial as a blank to account for background radioactivity. The samples were incubated for 5 min at room temperature, and their radiation signals were measured as counts per minute (CPM) using a Beckman-Coulter LS6500 scintillation counter. The CPM values of the blank were subtracted from each sample. The CPM values of KOH samples (accounting for newly synthesized DNA) were subtracted from the TCA-only samples (accounting for total nucleic acids) to calculate the CPM for newly synthesized RNA. The CPM values were finally normalized to $OD_{600}$ of each sample measured at the beginning of treatment.

Further, we confirmed that 0.02 µg/mL RIF and 1 µg/mL FDC inhibit RNA and DNA synthesis, respectively, to starvation levels. Briefly, overnight *S. aureus* cultures were centrifuged and resuspended in CA-MHB or PBS. For nucleic acid quantitation, 1.5 mL of each culture was dispensed into conical tubes containing 1.5 µCi $^3$H-uridine and treated with either RIF, FDC, or no inhibitor for 2 h. After 2 h, the samples were precipitated and analyzed as described above, and the CPM of each sample was determined.

**Timelapse microscopy.**

We performed timelapse microscopy to visualize morphological changes and *recA* expression in *S. aureus* persisters and non-persisters following FQ treatment using a method described previously (5). *S. aureus* HG003::pCM29-$P_{recA}$-*gfp*, the fluorescent reporter for *recA* transcription, was grown overnight for 18 h in 25 mL CA-MHB in a 250-mL baffled flask at 37 °C and 250 rpm with 10 µg/mL CAM. *S. aureus* HG003::$P_{recA}$-*recA-gfp*, encoding a translational fusion of RecA and GFP to mark DNA damage, was grown overnight for 18 h in 25 mL CA-MHB without any antibiotic. After 18 h, 2-mL cultures were dispensed into test tubes and treated with DMSO as a negative control, 5 µg/mL DLX, or 10 µg/mL CIP for 5 h at 37 °C and 250 rpm. After treatment, the cells were washed four times with 900 µL sterile PBS and finally resuspended in PBS. To starve cells after treatment, FQ-treated cells were incubated in PBS at 37 °C for 2 h before being inoculated onto agarose pads and imaged.

Meanwhile, during antibiotic treatment, we prepared agarose pads with CA-MHB and propidium iodide (PI; 1.6 μM; ThermoFisher) in Bioptechs chambers for imaging following a method described earlier (2). For imaging HG003::pCM29-$P_{recA}$-*gfp,* 10 μg/mL CAM was included in the pads for plasmid maintenance. After the agarose pads had dried for ~1 h, *S. aureus* cell suspensions were mixed thoroughly by vortexing to reduce clumping. The cells were diluted to $OD_{600}$ 0.05 in PBS, and 5 μL of cell suspension was spotted onto each agarose section marked by a 3D-printed divider. As soon as the drops dried, a 30-mm coverslip was placed onto the agarose pad, and the chamber was sealed.

A Zeiss Axiovert 200M microscope with a Plan-Apochromat 63x/1.40 Oil Ph3 M27 objective and environmental control was used for time-lapse image acquisition, and MetaMorph software Premier version 7.10.5 (Molecular Devices) was used to control the microscope and image acquisition. The phase contrast channel was utilized to locate and maintain the focal plane during multi-dimensional acquisition using MetaMorph's integrated autofocus algorithm. The samples were excited for fluorescence imaging of GFP (470/24) using Zeiss filter set 44 (beamsplitter FT 500, emission BP 530/50) and propidium iodide (Cy3; 555/15 nm) using Zeiss filter set 15 (Beamsplitter FT580, Emission LP590). Images were acquired on a pco.edge 4.2 bi sCMOS camera (6.5-mm pixel size). At least five different positions for each treatment condition were imaged every 10 min for 16 h. The images were processed and analyzed using ImageJ version 1.53 as described earlier (2).

Individual *S. aureus* pCM29-$P_{recA}$-*gfp* cells were observed during out-growth, and their fluorescence was quantified via ImageJ 1.54g using MicrobeJ version 5.13 (28). Cells were detected as circular shapes using the following parameters: area 50 $p^2$-max, length 1.1 p-max, and circularity 0-1. Masks placed by the software over individual cells were manually corrected when necessary. The persister and untreated cells were tracked until first division, and non-persisters were tracked for 25 frames or until they became PI-positive or lysed (i.e., when they die). We selected at least 25 untreated cells, 25 non-persisters from each drug treatment, and up to 25 persisters from each drug treatment for tracking using a random number generator to prevent bias. For DLX-treated populations, we tracked all 11 of the persisters we captured. Then, we determined the GFP signal of these cells.

For the translational DSB reporter (RecA-GFP) strain, analysis was performed using Fiji 1.54p with TrackMate v8.1.6 (29, 30). Cells were detected using the LoG (Laplacian of Gaussian) detector with the following parameters: diameter of 10 pixels and a threshold of 4. Spots were manually added when cells were not detected by the software. Persister and non-persister cells were tracked for the following parameters: presence or absence of foci, focus formation before or after division, fate of the focus, and uptake of PI. Additionally, the number of living persister progeny, defined as cells that had not taken up PI or lysed, for the first three generations was tracked. We tracked up to 10 persisters and 10 non-persisters per frame across replicates. For DLX, we tracked 31 persisters and 150 non-persisters for the starved condition and 19 persisters and 299 non-persisters for the unstarved condition (populations that were inoculated onto the pads after treatment). For CIP, we tracked 144 persisters and 141 non-persisters for the starved condition and 19 persisters and 72 non-persisters for the unstarved condition (populations that were fed right away).

**Quantification of Intracellular antibiotic abundance.**

Intracellular FQ abundance was measured using a method described previously (31). *S. aureus* HG003, *recA*::Tn, and *rexB*::Tn strains were grown overnight (18 h) in 25 mL CA-MHB in 250 mL flasks. Subsequently, 2 mL of each culture was transferred into glass tubes and treated with DLX (5 μg/mL) or CIP (10 μg/mL) for 5 h at 37 °C. After 5 h, 400 μL aliquots were collected and layered onto 350 μL of a silicone oil mixture (AR20: Sigma High Temperature, 9:1) pre-cooled to −80 °C, followed by centrifugation at 13,000 × *g* for 2 min at 4 °C. This step traps extracellular molecules within the oil layer, enabling measurement of only intracellular contents. The supernatant was then removed, and the cell pellets were stored at −80 °C for further processing. The remaining culture was washed four times with PBS as described above. Then, 400 μL of culture was processed using the silicone oil method after 0, 2, or 4 h of incubation in PBS at 37 °C with shaking and stored at −80 °C.

The following day, pellets were resuspended in 200 µL LC-MS grade water (Fisher Scientific), followed by the addition of 800 µL methanol: acetonitrile (1:1) (Fisher Scientific, LC-MS grade). Cloxacillin (50 ng/mL; Fisher Scientific) was spiked into each sample as an internal control for mass spectrometry analysis. Cells were transferred to tubes (USA Scientific) containing glass beads (Biospec products) and lysed by vortexing at maximum speed twice for 45 s on a Vortex Genie 2 (Scientific Industries), with a 2-min incubation on ice between cycles. Lysates were centrifuged at 21,000 × *g* for 10 min at 4 °C. A total of 600 µL supernatant was collected and centrifuged again for 3 min to remove debris. Finally, 400 µL of the clarified lysate was transferred to a new microcentrifuge tube and analyzed by mass spectrometry.

A 400-µL portion of the extract was collected into a pre-weighed 5-mL plastic tube (Eppendorf) and mixed with 3 times the volume of LC-MS Optima grade water to allow freezing at -80 °C. The frozen samples were dried using a lyophilizer (Millrock Technology, Inc). The dried extracts were reconstituted in 50 µL of acetonitrile: methanol: water (40:40:20) and centrifuged at 15,000 × *g* for 5 minutes. The supernatants were then transferred to LC-MS vials for analysis.

The samples were analyzed on a Synapt G2-Si QToF UPLC-MS/MS system (Waters Corp.) in positive ionization mode. Chromatographic separations were performed on an Acquity UPLC HSS T3 column (150 x 2.1 mm, 1.8 um particle size) (Waters Corp.) with the column temperature maintained at 40 °C. Water and acetonitrile containing 0.1% formic acid were used as solvents A and B, respectively. The mobile phase gradient with a flow rate at 0.3 mL/min was maintained as follows: 0 - 0.2 min: 5% B; 5 – 5.5 min: 90% B; and 6 – 8 min: 5% B. The injection volume was 10 µL. The data were acquired in the resolution mode. The scan ranges for MS and MSe acquisitions were 110 to 2000 Da and 50 to 2000 Da, respectively. A collision energy ramp of 5 to 40 V was applied for MSe acquisition. Leucine enkephalin (400 pg/µL), at an infusion flow rate of 10 µL/min, was used for real-time lockspray correction. Extracted ion chromatograms for the target compounds were generated, and the peaks were integrated using MassLynx 4.2 software (Waters Corp.)

Standards with 0, 10, 50, 100, 500, or 1000 ng/mL of DLX, CIP, or CLX were analyzed as described above to generate standard curves. The area under the curve (AUC) for a given sample was plotted as a function of known concentration using Microsoft Excel version 16.107.3. The y-intercept of each plot was set to 0, and a linear trendline was fit to each dataset. The equation for each compound's trendline was used to calculate the concentration of that compound in a given sample from the sample's AUC value for that compound. To account for volume loss that may have occurred during bead-beating, each sample's FQ concentration was normalized to its CLX concentration.

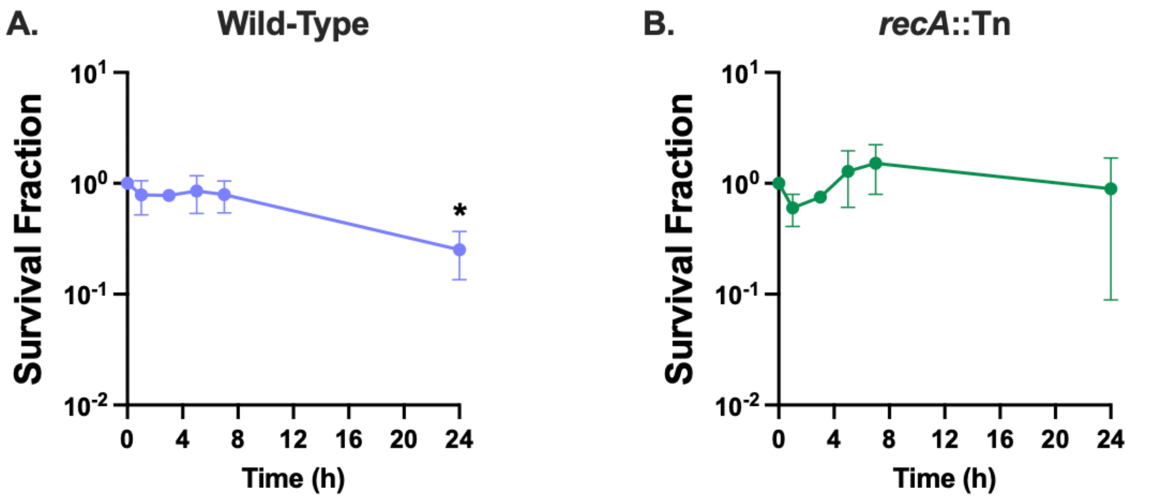


**Fig. S1.** *Cells are in stationary phase at the time of FQ treatment.* 18-h cultures of *S. aureus* (A) HG003 WT or (B) HG003 *recA*::Tn were incubated for an additional 24 h, and survival fractions were determined at designated timepoints. The cells did not proliferate during this time. At least three independent replicates were performed. *P* values were calculated by comparing log-transformed values at each later timepoint to 0 h using Dunnett's multiple comparisons test following analysis of variance (ANOVA). **P* < 0.05. Error bars denote SEM.

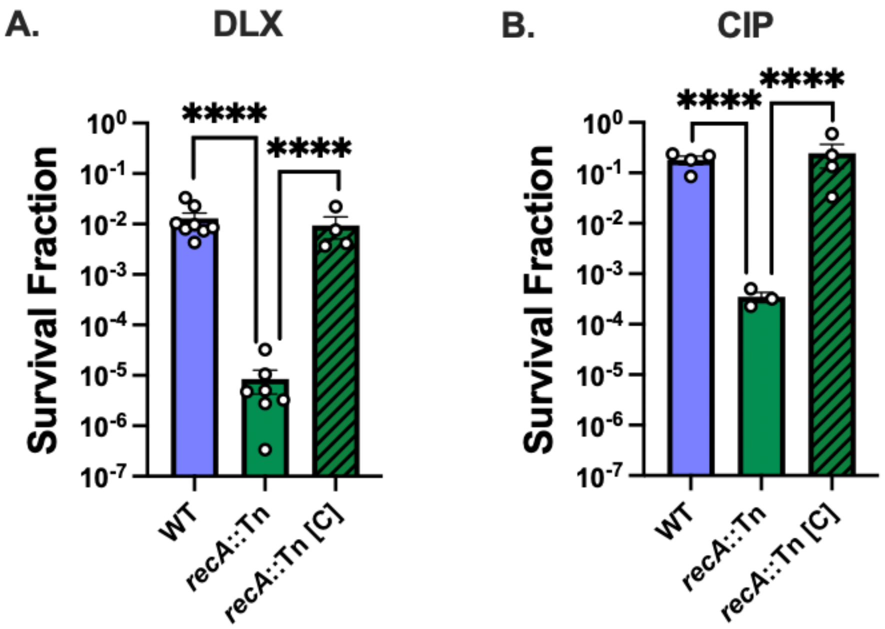


**Fig. S2.** *Loss of* recA *decreases* S. aureus *FQ survival.* (A-B) Stationary-phase cultures of WT, *recA*::Tn, and *recA*::Tn complemented with $P_{recA}$-*recA* (*recA*::Tn [C]) were treated for 5 h with (A) 5 µg/mL DLX or (B) 10 µg/mL CIP before survival was assessed. *P* values were calculated using Dunnett's multiple comparisons test following ANOVA to compare the log-transformed survival fraction of each strain to that of the other two strains treated with the same drug. At least three independent replicates were performed for each experiment. ****$P < 0.001$. Error bars denote SEM.

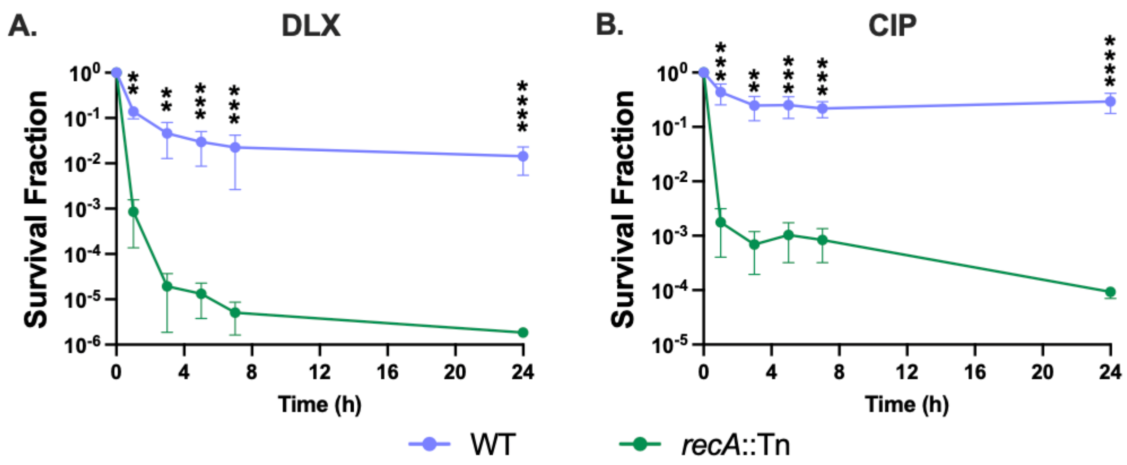


**Fig. S3.** *Cells surviving at 5 h of FQ treatment are persisters.* (A-B) Stationary-phase cultures of WT and *recA*::Tn were treated for 24 h with (A) 5 µg/mL DLX or (B) 10 µg/mL CIP, and survival was measured at indicated timepoints. WT and *recA*::Tn treated with either drug show a sharp decrease in survival fraction over the first 3 h, followed by a plateau indicative of a persister subpopulation. *P* values were calculated using *t*-tests to compare log-transformed survival fractions between WT and *recA*::Tn at each timepoint. At least three independent replicates were performed for each experiment. ***P* < 0.01, ****P* < 0.005, *****P* < 0.001. Error bars denote SEM.

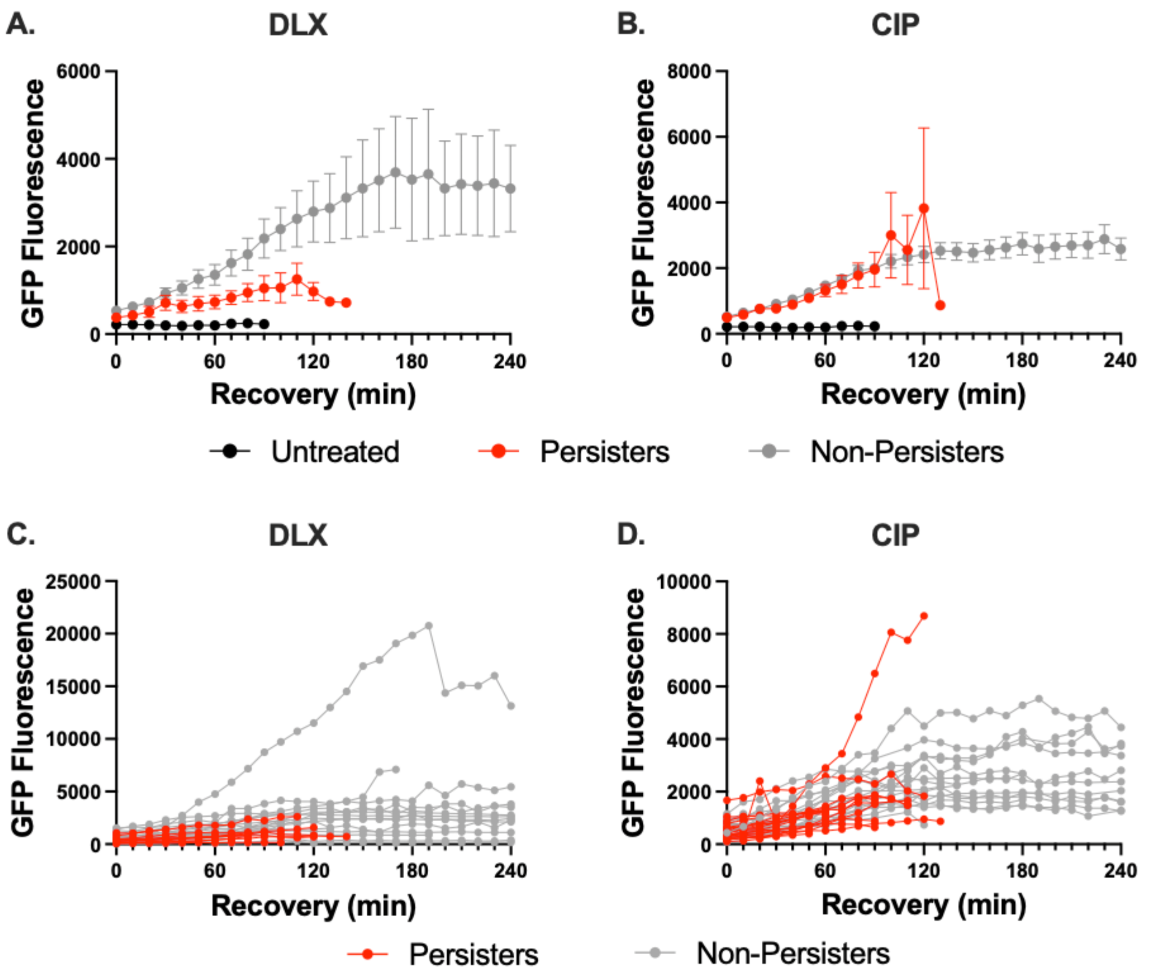


**Fig. S4.** *Both FQ persisters and non-persisters express* recA *during post-treatment recovery.* GFP signal of 26 untreated cells, 11 DLX persisters, 25 DLX non-persisters, 25 CIP persisters, and 25 CIP non-persisters was tracked via timelapse microscopy. Persisters were tracked until the point of division, and non-persisters were tracked for 240 min or until the point of death. Except for the 11 DLX persisters, all tracked cells were selected using a random number generator from cells imaged during at least two independent experiments. (A-B) Mean and (C-D) individual GFP fluorescence intensities are presented.

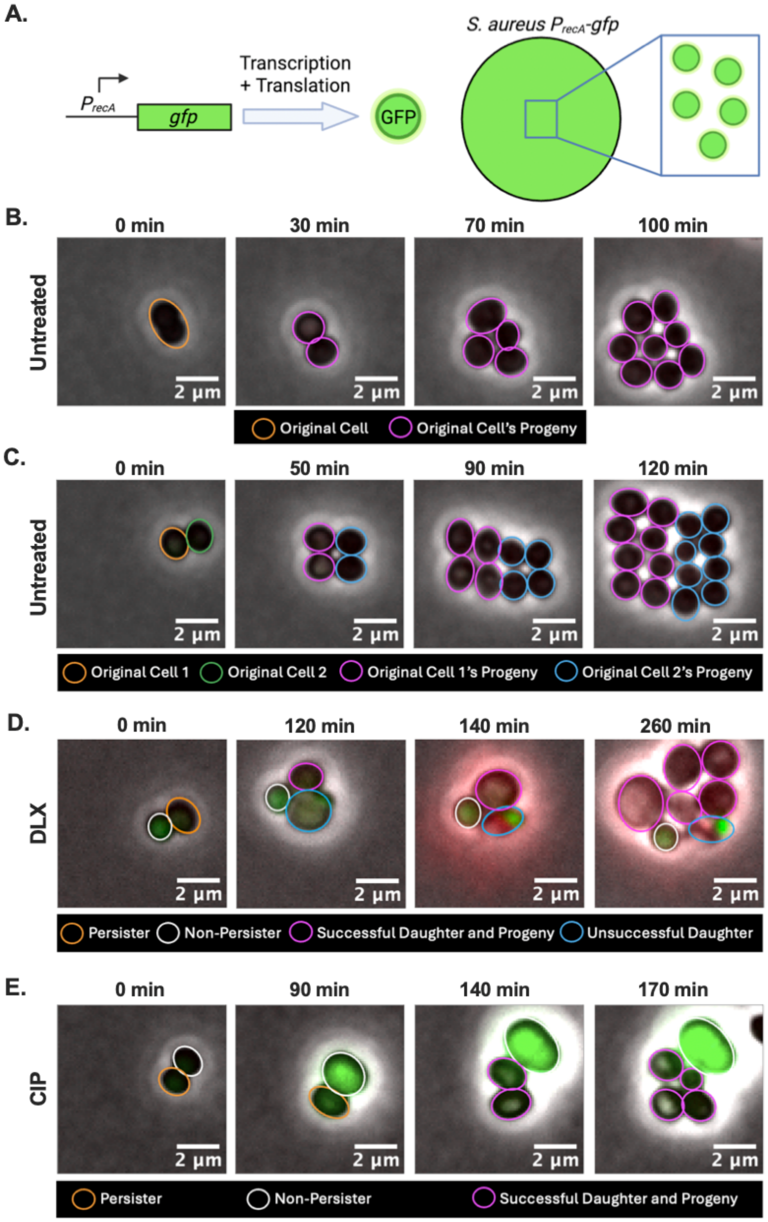


**Fig. S5.** *Comparison of* recA *expression and cell division between untreated and treated populations.* (A) Schematic illustrating *P*$_{recA}$*-gfp* construct in which GFP is expressed under the control of HG003's *recA* promoter. Created with BioRender. (B-E) Stationary-phase HG003::*P*$_{recA}$-*gfp* was monitored via timelapse microscopy on a Mueller-Hinton agarose pad after (B-C) no treatment, (D) DLX treatment, or (E) CIP treatment. (B-C) Each untreated cell present at the beginning of imaging divides into two cells, which continue to produce living progeny. No appreciable *gfp* expression (indicating *recA* induction) or PI staining (indicating loss of membrane integrity in dying/dead cells) is apparent in untreated cells. (D) The DLX-treated non-persister (white outline) turns green, indicating *recA* expression and DNA damage. The DLX persister (orange outline) divides into two daughters. The blue daughter expresses *recA,* fails to divide further, and turns PI-positive while the magenta divides normally. (E) The CIP-treated non-persister (white outline) swells and shows high *recA* expression. The CIP persister (orange outline) shows modest *recA* expression, divides, and produces two daughters that both divide. Microscopy images are representative of two independent replicates.

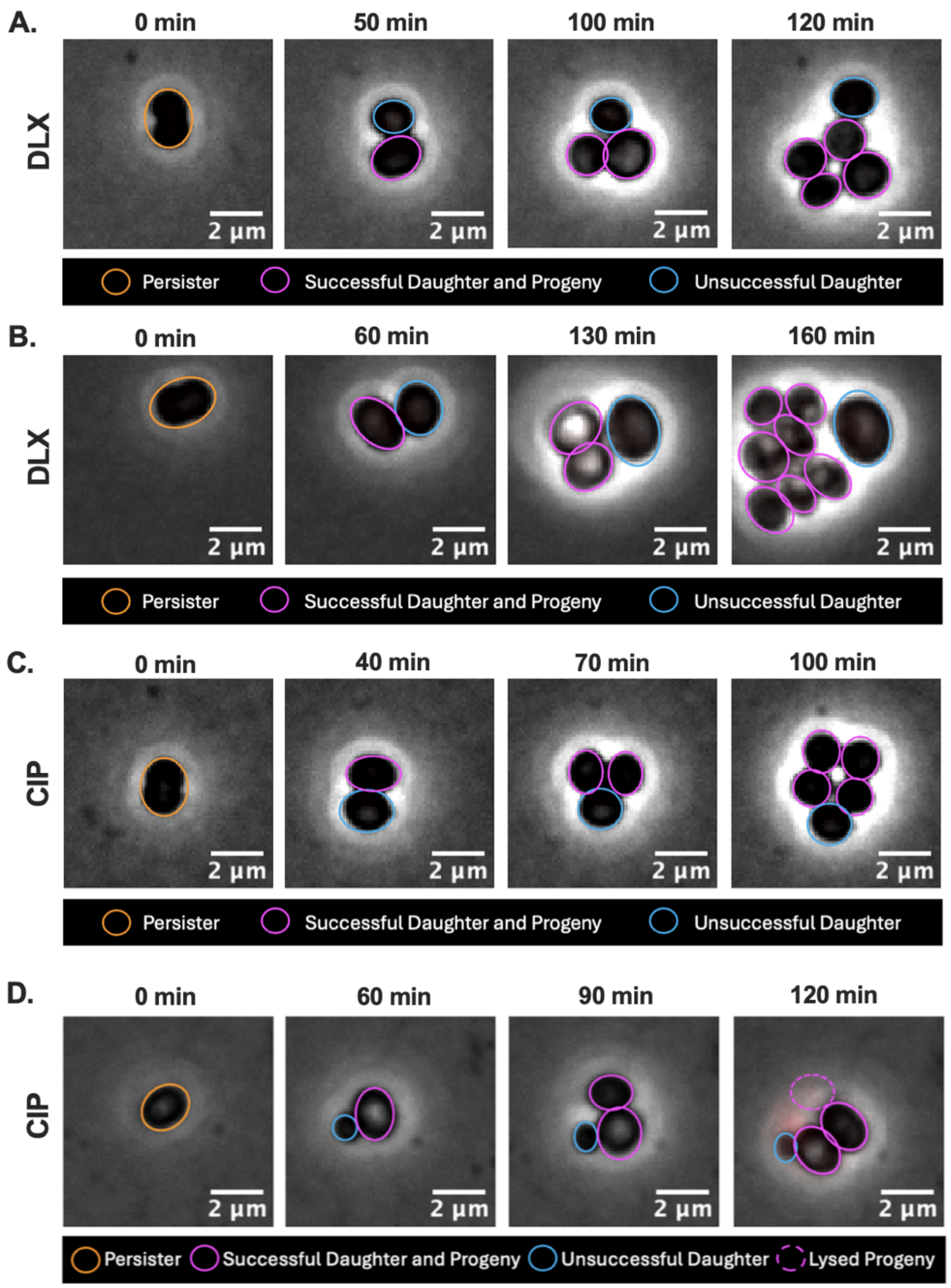


**Fig. S6.** *Stalling of persisters daughters is not an artefact of reporter constructs.* Reporter-free HG003 was treated for 5 h with (A-B) DLX or (C-D) CIP then observed via timelapse microscopy during recovery on Mueller-Hinton agarose. Similar to HG003::$P_{recA}$-*gfp,* the reporter-free FQ-treated persisters often produced one successful daughter, which continued to divide and produce progeny, and one unsuccessful daughter, which failed to divide again. Images are representative of two independent experiments.

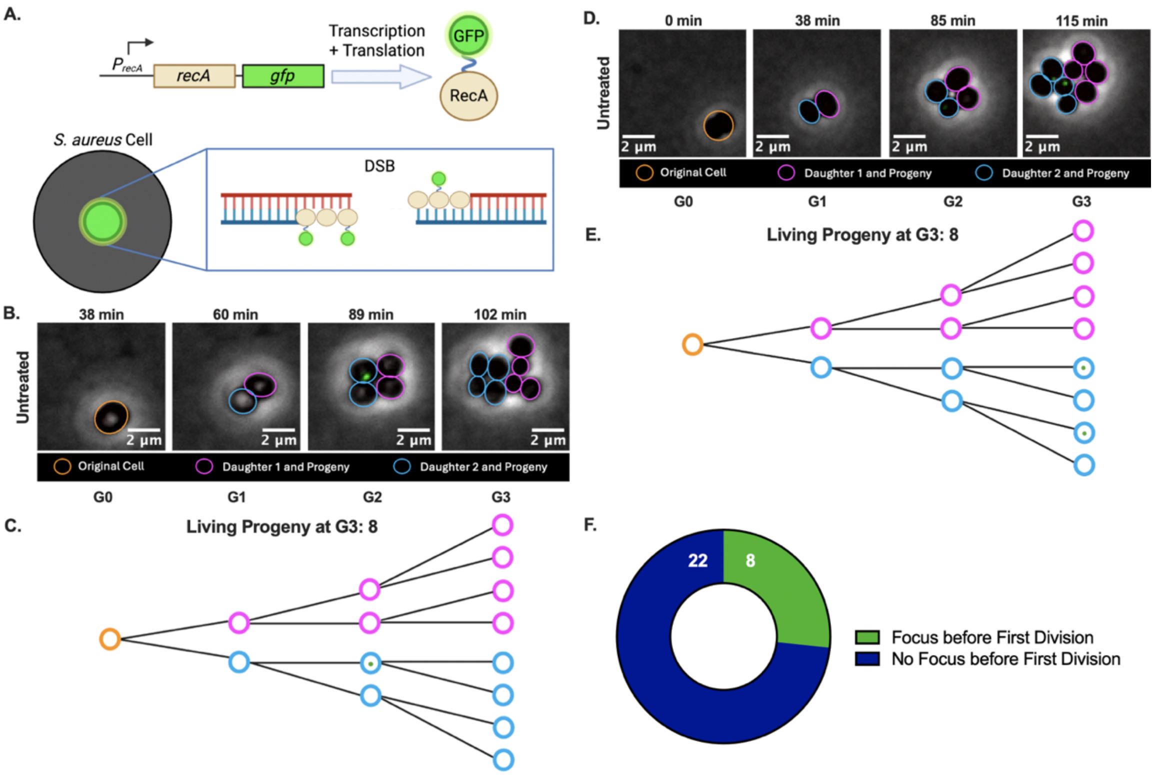


**Fig. S7.** *Untreated cells divide normally and usually do not produce RecA-GFP foci before division.* (A) Schematic illustrating $P_{recA}$-*recA-gfp* construct in which RecA and GFP are expressed under the control of HG003's *recA* promoter and connected by a flexible linker. RecA-GFP proteins bind to single-stranded DNA that arises during DSB processing, producing fluorescent foci marking DSBs. Non-GFP-tagged RecA molecules are also shown since this strain also contains the native *recA* gene. Created with BioRender. (B-F) Untreated HG003::$P_{recA}$-*recA-gfp* were monitored via timelapse microscopy on Mueller-Hinton agarose pads. (B) An untreated cell (G0, orange outline) divides into two daughters at G1 (blue and magenta). Both daughters continue to divide, and one blue progeny has a focus at G2. (C, E) Schematics illustrating the division events observable in (B) and (D), respectively, as well as the number of living progeny at the end of G3. Circles represent cells, and green dots represent RecA-GFP foci. Created with BioRender. (D) An untreated cell (G0, orange outline) divides into two daughters (G1, blue and magenta). Each daughter cell continues to divide, resulting in four and eight cells at G2 and G3, respectively. Two G3 cells resulting from the blue daughter have foci. (F) Thirty randomly selected untreated single cells across three independent replicates were analyzed, and the proportion that had a RecA-GFP focus at any point before their first division was calculated.

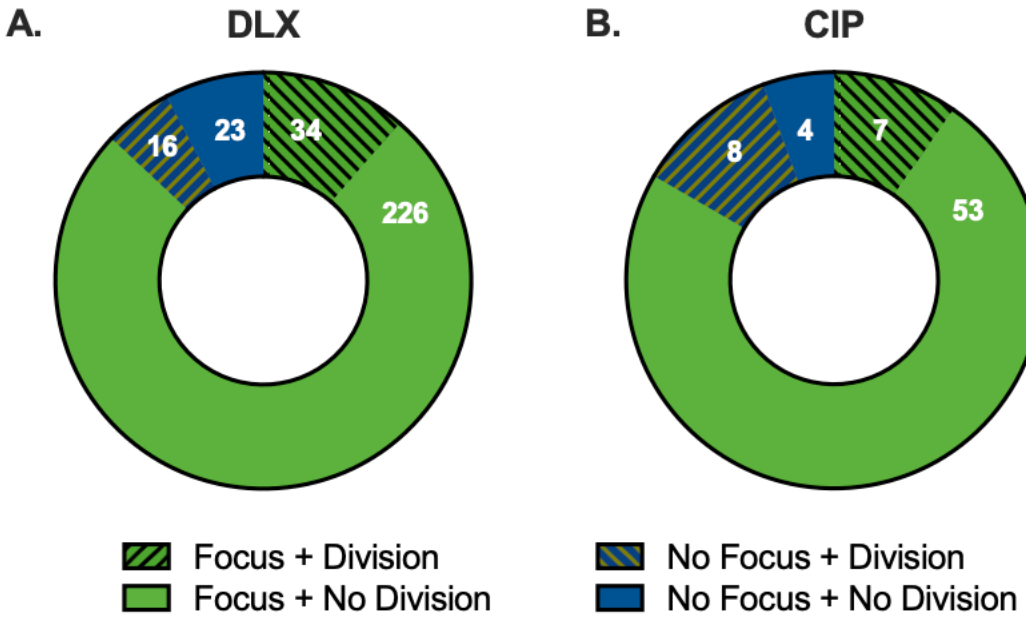


**Fig. S8.** *Most FQ-treated non-persisters have RecA-GFP foci and do not divide.* Stationary-phase cultures of HG003::$P_{recA}$-*recA-gfp* were treated for 5 h with DLX or CIP then observed via timelapse microscopy. From at least three independent experiments, 299 DLX-treated non-persisters and 72 CIP-treated non-persisters were randomly selected for analysis. The proportion of (A) DLX-treated and (B) CIP-treated non-persisters that had a RecA-GFP focus or no focus, and divided or not, was calculated. Although some of these cells divided, they were not considered persisters because both of their daughters stalled, resulting in failure to produce a microcolony.

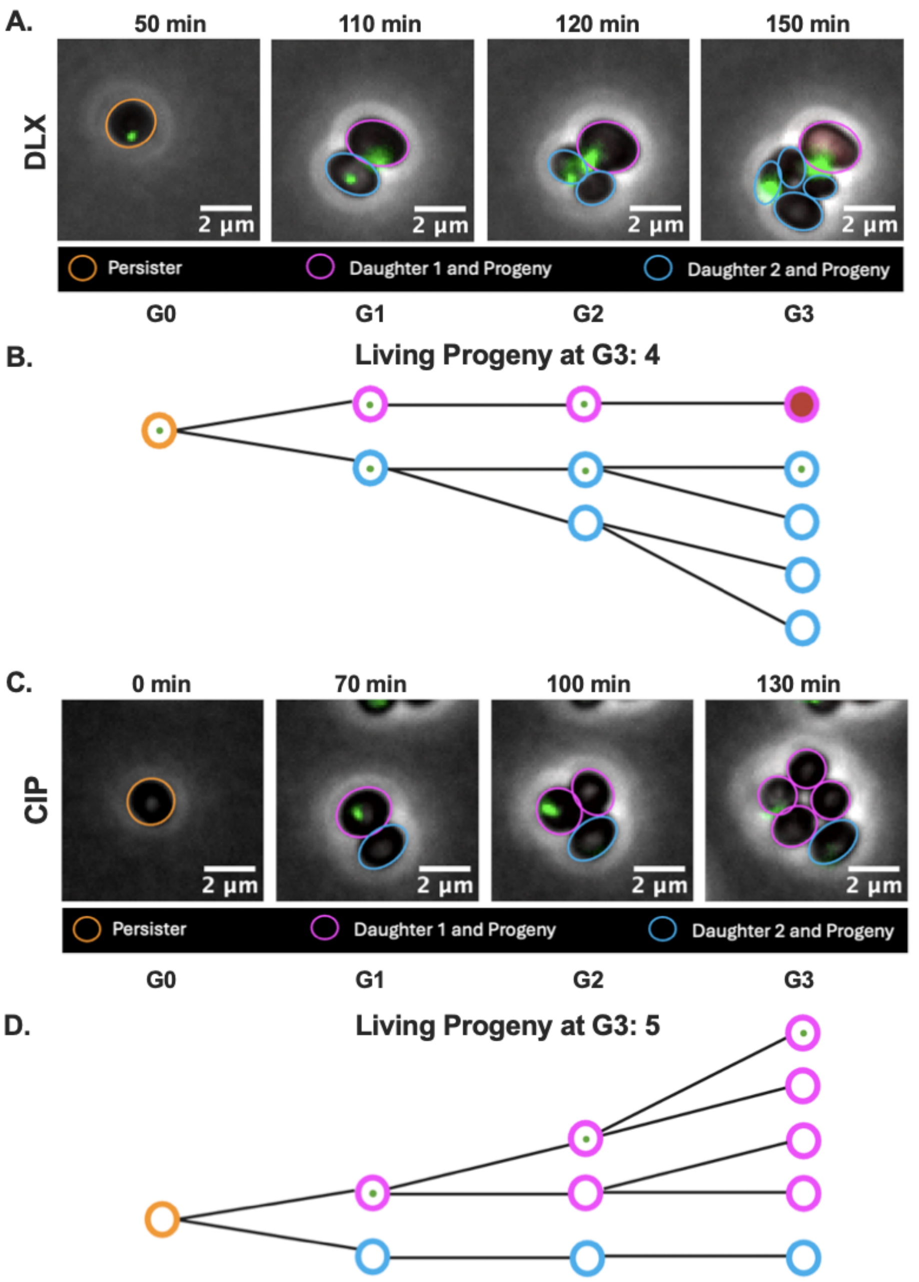


**Fig. S9.** *Persisters frequently produce only one successful daughter.* Stationary-phase HG003::*P*$_{recA}$-*recA-gfp* was treated for 5 h with (A-B) DLX or (C-D) CIP then monitored via timelapse microscopy. (A) A persister (orange outline, G0) has a RecA-GFP focus then divides into two cells (blue and magenta, G1), each of which has a focus. The blue daughter proliferates, whereas the magenta daughter stalls and eventually takes up PI, indicating loss of membrane integrity and death. (B, D) Schematics illustrating the division events observable in (A) and (C), respectively, as well as the number of living progeny (defined as cells that have not taken up PI or lysed) at the end of G3. Circles represent cells, green dots represent RecA-GFP foci, and red shading indicates PI staining. Created with BioRender. (C) A persister (orange outline, G0) divides into two cells, (blue and magenta, G1). The magenta cell has a focus yet divides, producing four progenies by G3. The blue cell stalls but remains alive. Microscopy images are representative of at least three independent imaging experiments.

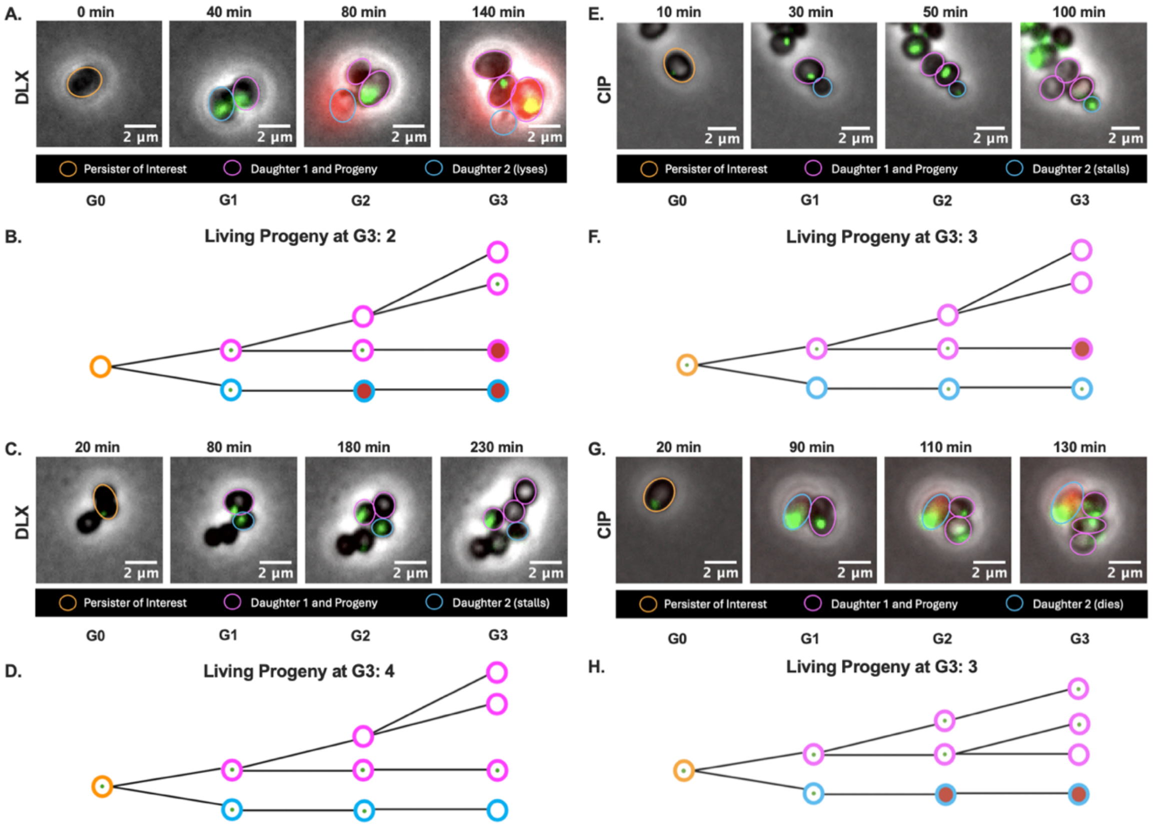


**Fig. S10.** *FQ persister progeny suffer consequences of treatment for several generations.* Stationary-phase HG003::*P*$_{recA}$-*recA-gfp* was treated for 5 h with (A-D) DLX or (E-H) CIP, then monitored via timelapse microscopy. (A) A persister (orange outline, G0) has no RecA-GFP focus and then divides into two cells (blue and magenta, G1), each of which has a focus. Daughter 1 (magenta outline) divides, and its G2 progeny that inherits the focus takes up PI, indicating loss of membrane integrity. The other G2 progeny divides, and one of its G3 progeny has a focus. Daughter 2 (blue outline) lyses. (B, D, F, H) Schematics illustrating the division events observable in (A), (C), (E), and (G), respectively, as well as the number of living progeny at the end of G3. Circles represent cells, green dots represent RecA-GFP foci, and red shading indicates PI staining. Created with BioRender. (C) A persister (orange outline, G0) has a focus and divides into two daughters (blue and magenta, G1), each of which has a focus. Daughter 1 (magenta outline) divides, and its G2 progeny that inherits the focus stalls. Its other G2 progeny divides successfully. Daughter 2 (blue outline) never divides. (E) A persister (orange outline, G0) has a focus and divides into two daughters, (blue and magenta, G1). Daughter 1 (magenta outline) has a focus yet divides. Its G2 progeny that inherits the focus stalls and later takes up PI, but the other G2 progeny divides successfully. Daughter 2 (blue outline) forms a focus and never divides. (G) A persister (orange outline, G0) has a focus and divides into two daughters, (blue and magenta, G1), each of which has a focus. Daughter 1 (magenta outline) divides into two cells that both have a focus. One of these G2 progeny stalls while the other divides, producing one G3 progeny with a focus and one with no focus. Daughter 2 (blue outline) takes up PI. Microscopy images are representative of at least three independent imaging experiments.

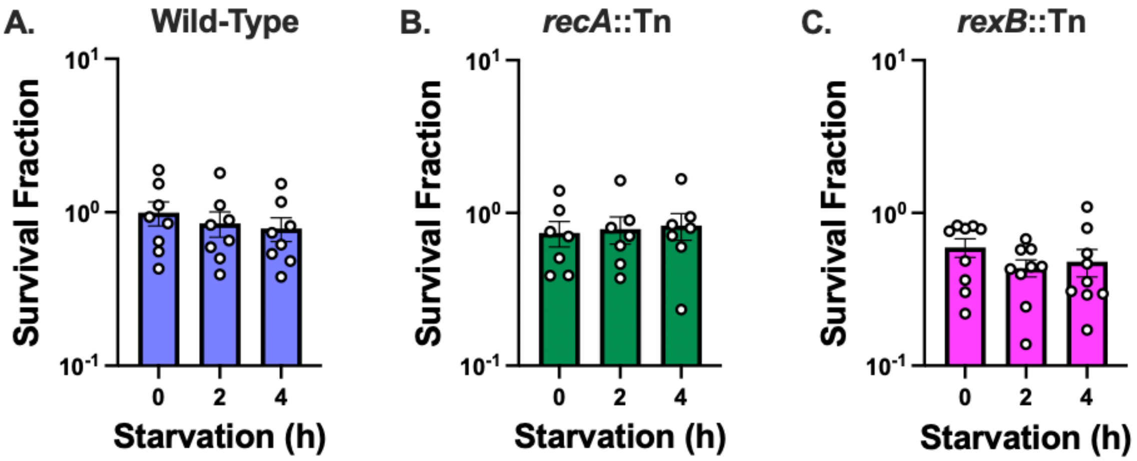


**Fig. S11.** *Effects of starvation on* S. aureus *culturability.* 18-h cultures of (A) WT, (B) *recA*::Tn, and (C) *rexB*::Tn were treated with DMSO (negative control) for 5 h then starved for 0, 2, or 4 h following treatment before survival was determined. At least seven independent replicates were performed for each experiment. *P* values were calculated using Dunnett's multiple comparisons test following ANOVA to compare log-transformed survival fractions at each timepoint to 0 h. No statistically significant differences ($P < 0.05$) were detected, indicating that 4 h of starvation did not affect the culturability of untreated cells. Error bars denote SEM.

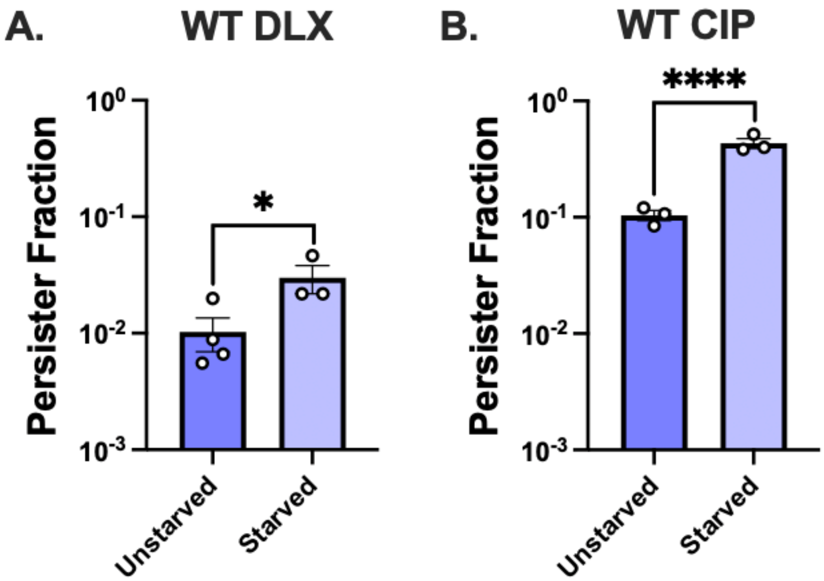


**Fig. S12.** *Timelapse microscopy confirms that post-treatment starvation increases the proportion of persisters in FQ-treated populations.* Stationary-phase cultures of HG003::$P_{recA}$-*recA-gfp* were treated for 5 h with (A) DLX or (B) CIP, starved or unstarved for 2 h, then observed via timelapse microscopy in at least three independent experiments. To determine the persister fraction, we divided the number of persisters (cells that gave rise to a microcolony) by the total number of cells (persisters and non-persisters) in each frame. *P* values were calculated using *t*-tests assuming equal variances to compare the log-transformed persister fractions between unstarved and starved populations. *$P < 0.05$, ****$P < 0.001$. Error bars denote SEM.

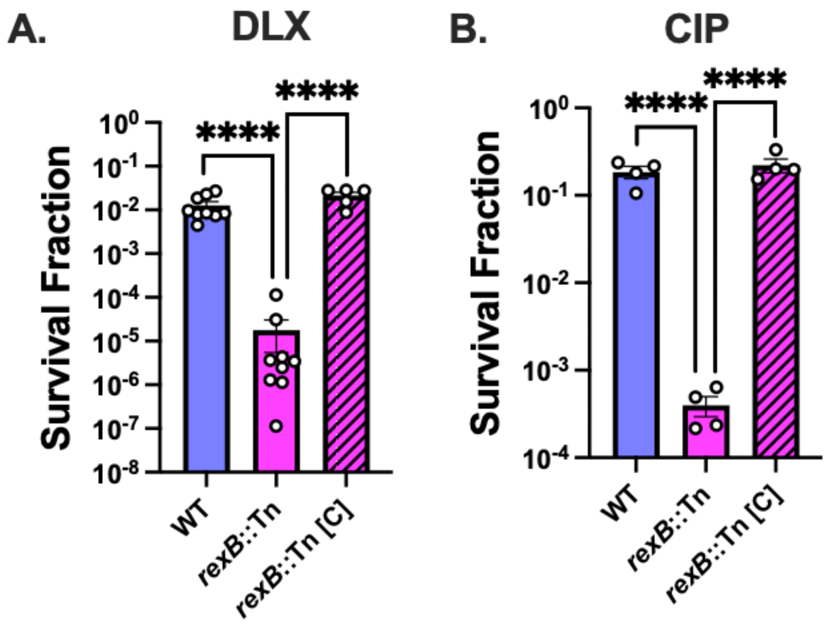


**Fig. S13.** *Disruption of* rexB *decreases FQ persistence.* (A-B) Stationary-phase WT, *rexB*::Tn, and *rexB*::Tn [C], where *rexB* is complemented on the chromosome, were treated with (A) 5 µg/mL DLX or (B) 10 µg/mL CIP for 5 h before survival was determined. At least three independent replicates were performed. *P* values were calculated using Dunnett's multiple comparisons test following ANOVA to compare log-transformed survival fractions of each strain treated with a given FQ. ****$P$ < 0.001. Error bars denote SEM.

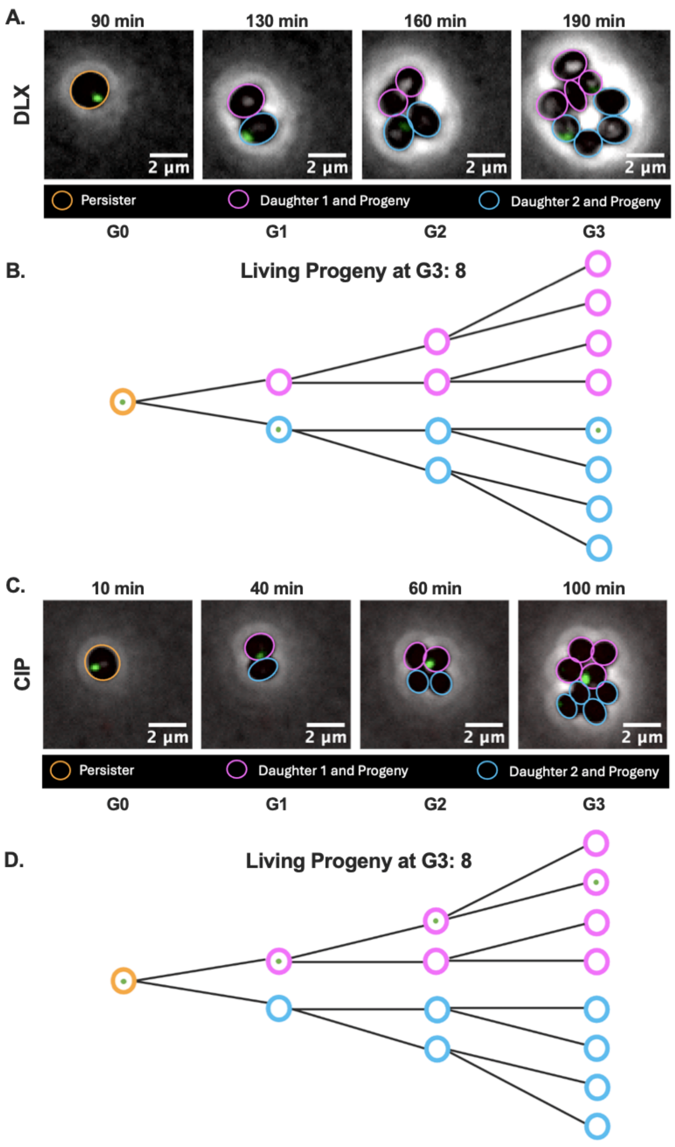


**Fig. S14.** *Starved FQ-persisters and their progeny frequently divide normally.* Stationary-phase cultures of *S. aureus* HG003::$P_{recA}$-*recA-gfp* were observed after treatment with DLX, CIP, or a negative control followed by starvation or immediate feeding. (A) A starved DLX persister (orange outline, G0) has a focus and divides into two cells (blue and magenta, G1). Although the daughter with the blue outline inherits the focus, both daughters divide normally, producing 4 total progeny at G2 and 8 at G3, similar to untreated cells (compare with Fig. S7B-F). One blue progeny at G3 has a focus. (B, D) Schematics illustrating the division events observable in (A) and (C), respectively, as well as the number of living progeny at the end of G3. Circles represent cells, and green dots represent RecA-GFP foci. Created with BioRender. (C) A starved CIP persister (orange outline, G0) has a focus and divides into two cells (blue and magenta, G1), which further divide normally. The cell with the magenta outline at G1 has a focus that is inherited by one of its progeny in each generation. The frequent lack of growth stalling and death among starved persister progeny suggests that post-treatment starvation may limit DNA damage not only in persisters but also in their progeny. Images are representative of at least three independent microscopy experiments.

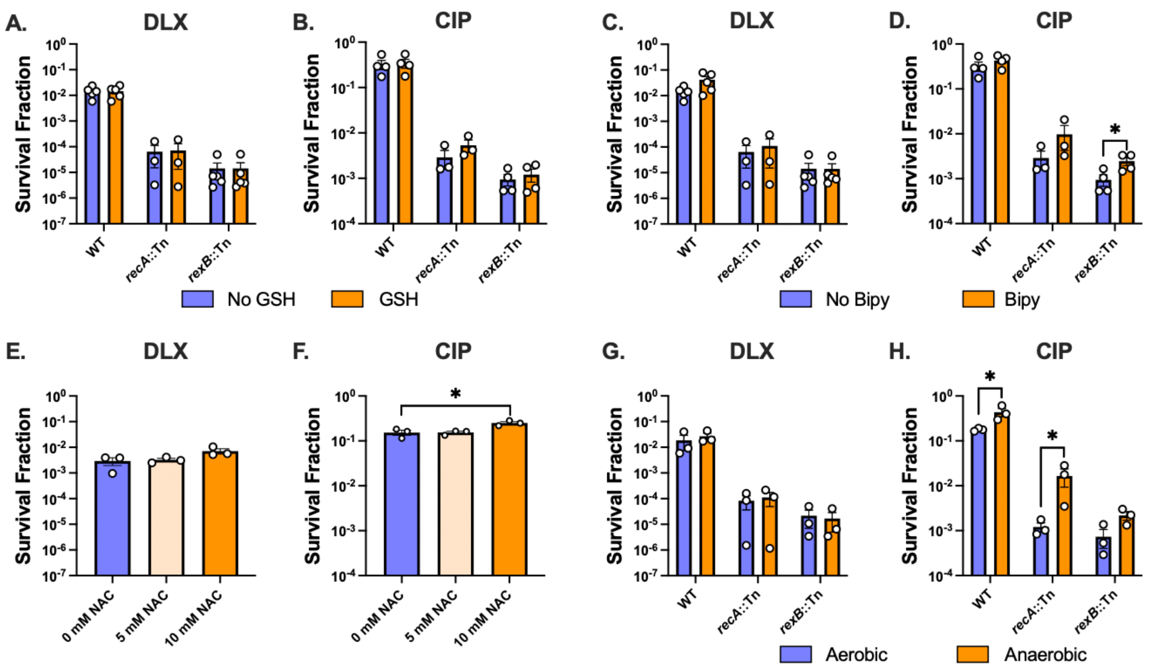


**Fig. S15.** *Limiting oxidative stress during recovery may improve CIP, but not DLX, survival.* Indicated *S. aureus* strains were treated with (A, C, E, G) 5 µg/mL DLX or (B, D, F, H) 10 µg/mL CIP then recovered (A-B) aerobically with or without GSH, (C-D) aerobically with or without Bipy, (E-F) aerobically with or without NAC, or (G-H) anaerobically. (A-D, G-H) *P* values were calculated using *t*-tests to compare log-transformed survival fractions of each strain treated with a given FQ then recovered (A-D) with vs. without a given antioxidant or (G-H) with vs. without oxygenation. (E-F) *P* values were calculated using Dunnett's multiple comparisons test following ANOVA to compare log-transformed survival fractions at each dose of NAC to no NAC. At least three independent replicates were performed. $^{*}P < 0.05$. Only significant comparisons are shown. Error bars denote SEM.

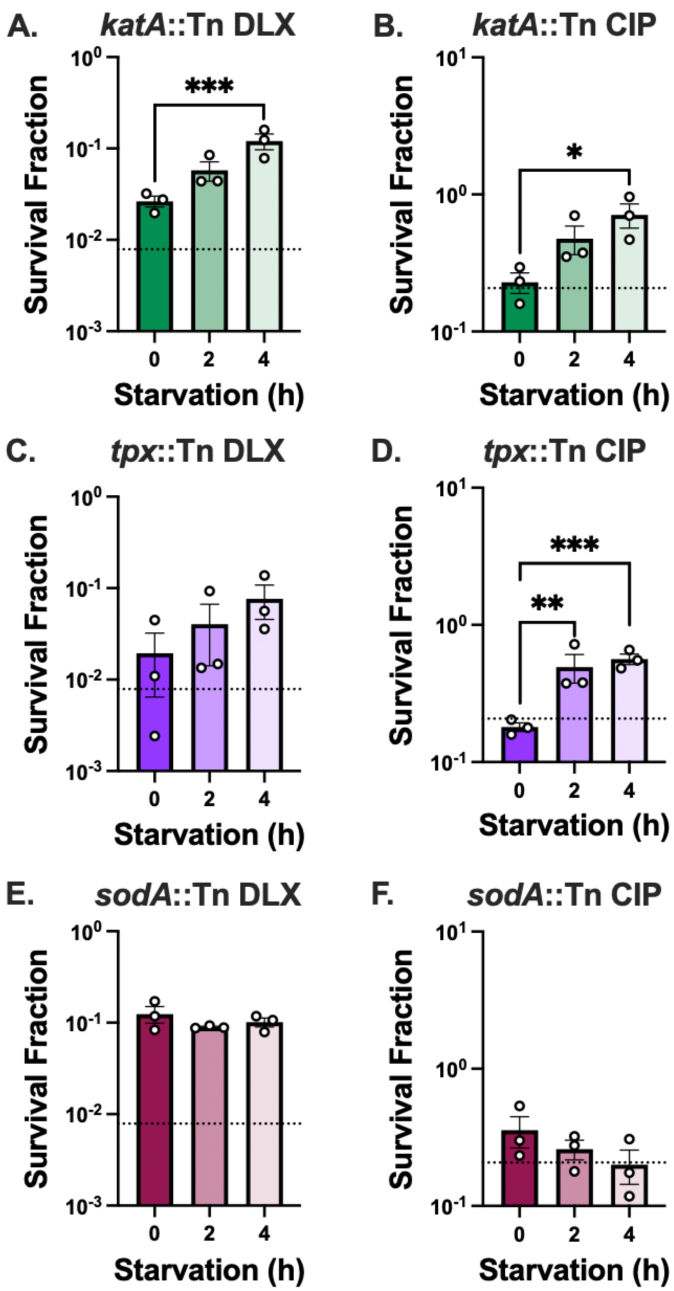


**Fig. S16.** *Starvation after FQ treatment increases survival of mutants deficient in ROS detoxification.* Stationary-phase *S. aureus* (A-B) *katA*::Tn, (C-D) *tpx*::Tn, and (E-F) *sodA*::Tn were treated with 5 µg/mL DLX or 10 µg/mL CIP for 5 h. After treatment, the cells were starved for 0, 2, or 4 h before being plated onto MHA for CFU enumeration. Dotted lines indicate the mean survival of non-starved WT cells treated with a given FQ. Three independent replicates were performed for each experiment. *P* values were calculated using Dunnett's multiple comparisons test following ANOVA to compare log-transformed survival fractions at each timepoint to 0 h. **P* < 0.05, ***P* < 0.01, ****P* < 0.005. Error bars denote SEM.

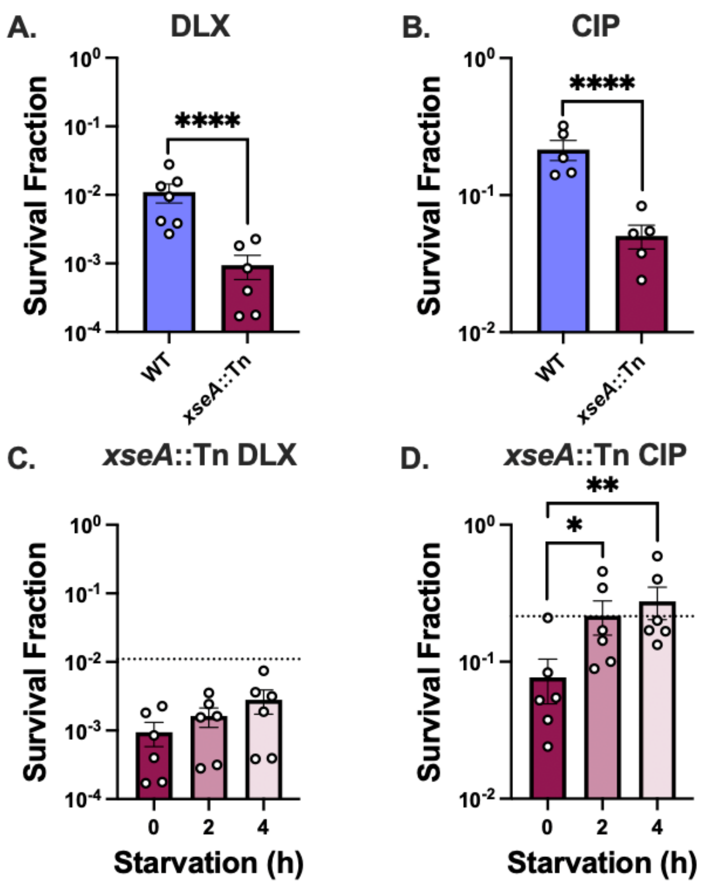


**Fig. S17.** *Loss of* xseA *decreases FQ survival but does not prevent starvation-induced survival increase.* (A-B) Stationary-phase *S. aureus* HG003 WT and *xseA*::Tn were treated for 5 h with (A) 5 µg/mL DLX or (B) 10 µg/mL CIP then recovered on MHA. (C-D) DLX- or CIP-treated *xseA*::Tn were fed immediately or starved for 2 or 4 h before being plated onto MHA for CFU enumeration. Dotted lines indicate the mean survival of unstarved WT cells treated with a given FQ. Three independent replicates were performed for each experiment. *P* values were calculated using (A-B) *t*-tests assuming equal variance to compare the log-transformed survival fraction of WT to *xseA*::Tn or (C-D) Dunnett's multiple comparisons test following ANOVA to compare the log-transformed survival fractions at each timepoint to 0 h. *$P < 0.05$, **$P < 0.01$, ****$P < 0.001$. Error bars denote SEM.

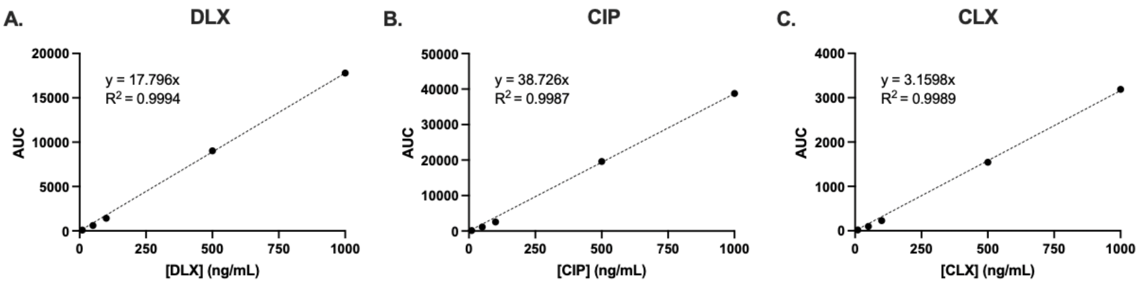


**Fig. S18.** *Representative standard curves of DLX, CIP, and CLX obtained using LC-MS/MS.* The area under the curve (AUC) resulting from LC-MS/MS analysis of a known concentration of each drug was plotted, and a linear trendline was fit to the data. The y-intercepts were set to 0, and the equations and $R^2$ values were obtained. Curves shown are representative of three independent replicates.

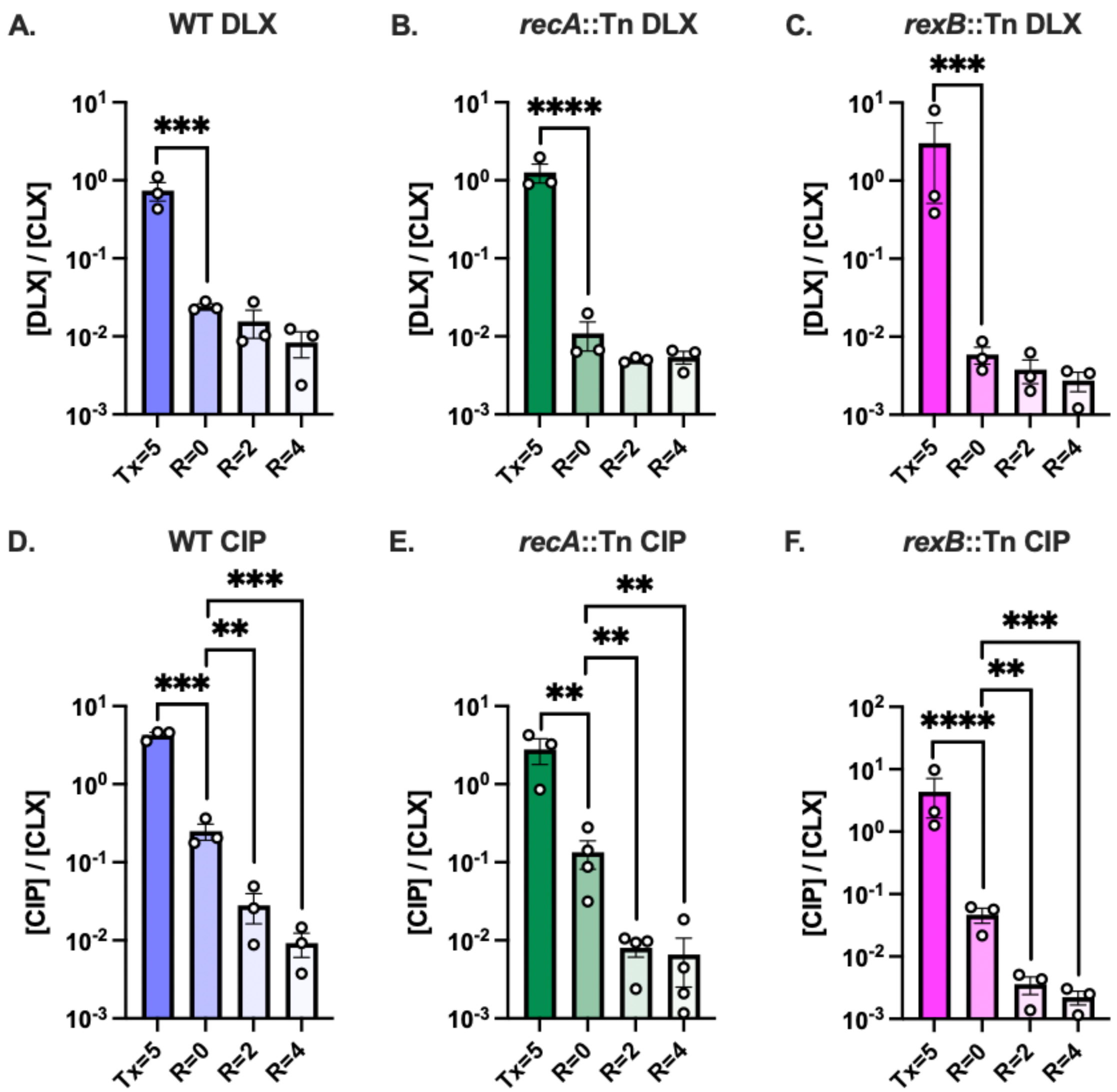


**Fig. S19.** *Post-treatment starvation may allow FQ expulsion before nutrient replenishment.* Stationary-phase *S. aureus* HG003 WT, *recA*::Tn, and *rexB*::Tn were treated for 5 h with (A-C) DLX or (D-F) CIP. Samples were removed for lysing and subsequent LC-MS/MS analysis immediately after treatment but before washing (Tx=5), immediately after washing (R=0), after 2 h of starvation in PBS (R=2), and after 4 h of starvation in PBS (R=4). After all extracellular drug was removed from each sample, cloxacillin (CLX) was spiked in for normalization immediately before the cells were lysed. At least three independent replicates were performed. *P* values were calculated using Dunnett's multiple comparisons test following ANOVA to compare the log-transformed [FQ]/[CLX] ratio at each timepoint to R=0. ***P* < 0.01, ****P* < 0.005, *****P* < 0.001. Only significant comparisons are shown. Error bars denote SEM.

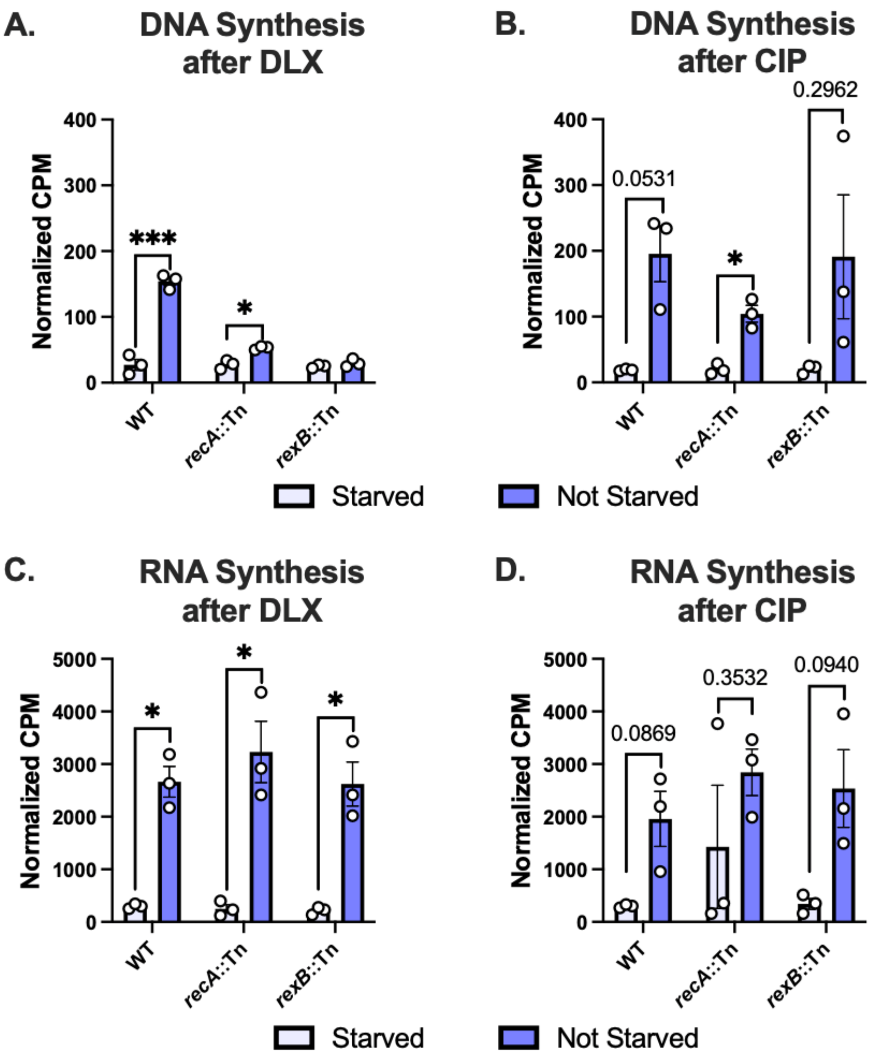


**Fig. S20.** *Starvation inhibits the resumption of nucleic acid synthesis after FQ treatment. S. aureus* WT, *recA*::Tn, or *rexB*::Tn were treated for 5 h with (A, C) 5 µg/mL DLX or (B, D) 10 µg/mL CIP then recovered for 2 h in the presence of $^3$H-uridine in PBS (starved) or CA-MHB (not starved) before incorporation of $^3$H-uridine into (A-B) DNA and (C-D) RNA was measured. *P* values were calculated using *t*-tests to compare normalized counts per minute (CPM) values for DNA or RNA of a given strain after recovery under starved vs. non-starved conditions. *$P < 0.05$, ***$P < 0.005$. Error bars denote SEM.

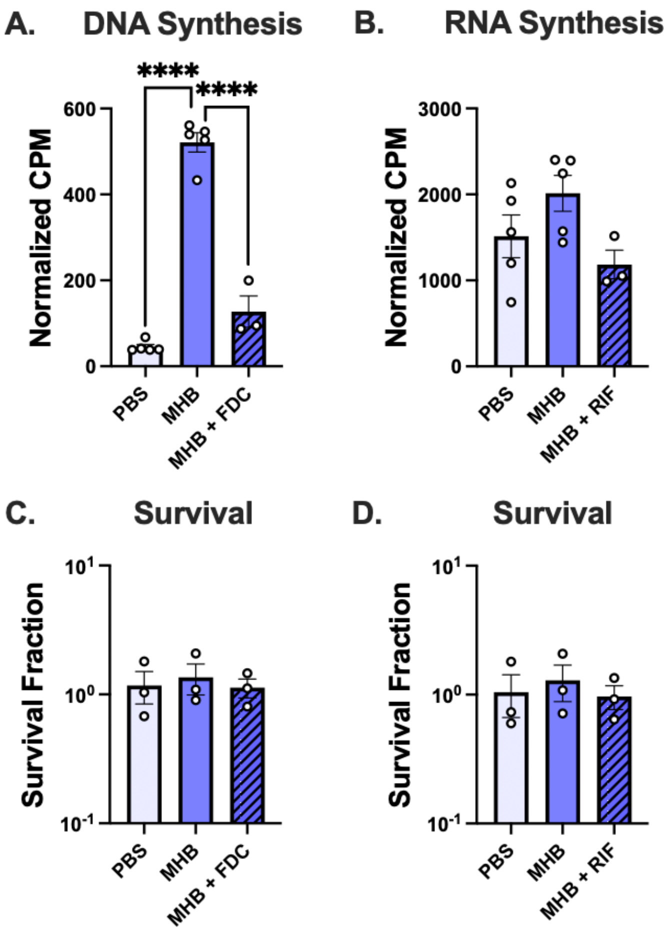


**Fig. S21.** *Doses of nucleic acid synthesis inhibitors used to inhibit DNA/RNA synthesis without killing* S. aureus. Stationary-phase *S. aureus* HG003 cultures were incubated for 2 h in PBS, MHB, (A, C) MHB + 1 µg/mL FDC or (B, D) MHB + 0.02 µg/mL RIF. (A, B) Cells were incubated with $^{3}$H-uridine with or without inhibitors before (A) DNA or (B) RNA synthesis was measured. (C, D) Survival of cells in each treatment condition was also determined. (A-B) *P* values were calculated using Dunnett's multiple comparisons test following ANOVA to compare normalized CPM values of each condition to MHB. ****$P$ < 0.001. Only significant comparisons are shown.

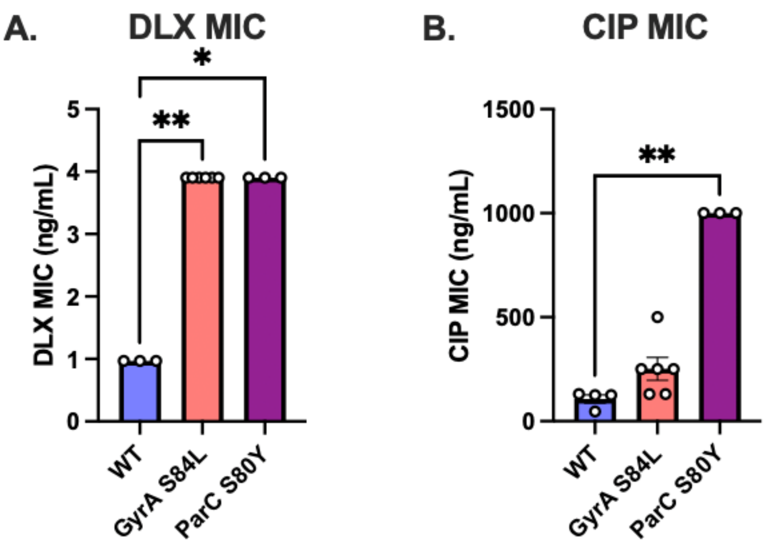


**Fig. S22.** *Effect of topoisomerase mutations on FQ MIC.* (A-B) Exponential-phase cultures of WT, GyrA S84L, and ParC S80Y were exposed to a range of (A) DLX or (B) CIP doses to determine MIC following CLSI guidelines. *P* values were calculated using Dunn's multiple comparisons test following the Kruskal-Wallis test to compare each mutant to WT. **P* < 0.05*,* ***P* < 0.01. At least three independent replicates were performed. Error bars denote SEM.

### Tables

#### **Table S1.** *Strains used in this study.*

| **Strain** | **Description** | **Source** |
|---|---|---|
| *S. aureus* HG003 | Wild-type parent strain | Dr. Brian Conlon (1) |
| *S. aureus* HG003 *recA*::Tn | *recA* transposon mutant | This study |
| *S. aureus* HG003 pBK123 | Plasmid with cadmium-inducible promoter ($P_{cad}$) | This study |
| *S. aureus* HG003 pBK123::$P_{cad}$-*recA* | Plasmid carrying *recA* under the control of the cadmium-inducible promoter | This study |
| *S. aureus HG003 PrecA-recA-gfp* | Genomic insertion of sequence encoding RecA-GFP translational fusion | This study |
| *S. aureus* HG003 *recA*::Tn [C] | *recA* transposon mutant strain with genomic complementation of $P_{recA}$-*recA* | This study |
| *S. aureus* HG003 *rexB*::Tn | *rexB* transposon mutant | This study |
| *S. aureus* HG003 *rexB*::Tn [C] | *rexB* transposon mutant strain with genomic complementation of $P_{rexB}$-*rexB* | This study |
| *S. aureus* HG003 *xseA*::Tn | *xseA* transposon mutant | This study |
| *S. aureus* HG003 *katA*::Tn | *katA* transposon mutant | This study |
| *S. aureus* HG003 *sodA*::Tn | *sodA* transposon mutant | This study |
| *S. aureus* HG003 *tpx*::Tn | *tpx* transposon mutant | This study |
| *S. aureus* JE2 | FQ-resistant strain | BEI Resources |
| *S. aureus* JE2 *recA*::Tn | *recA* transposon mutant | BEI Resources |
| *S. aureus* JE2 *rexB*::Tn | *rexB* transposon mutant | BEI Resources |
| *S. aureus* JE2 *xseA*::Tn | *xseA* transposon mutant | BEI Resources |
| *S. aureus JE2 katA*::Tn | *katA* transposon mutant | BEI Resources |
| *S. aureus JE2 sodA*::Tn | *sodA* transposon mutant | BEI Resources |
| *S. aureus JE2 tpx*::Tn | *tpx* transposon mutant | BEI Resources |
| *S. aureus* RN4220 | Restriction-negative cloning strain | Dr. Richard Novick (14) |
| *S. aureus* RN9011 | RN4220 containing pRN7023, which carries SaPI1 integrase | Dr. Francis Alonzo III (8-9) |
| *S. aureus* HG003::*gyrA*-S84L | HG003 containing chromosomal *gyrA*-S84L mutation | This study |
| *S. aureus* HG003::*parC*-S80Y | HG003 containing chromosomal *parC*-S80Y mutation | This study |

**Table S2.** *Plasmids used in this study.*

| Plasmid | Description | Resistance Marker | Source |
|---|---|---|---|
| pBK123 | Plasmid with cadmium-inducible promoter ($P_{cad}$) | Chloramphenicol | Dr. Paul Fey (10-11) |
| pCM29 | Plasmid with *gfp* under the control of the constitutive promoter *sarA* P1 | Chloramphenicol | Dr. Alexander Horswill (15) |
| pJC1111 | SaPI1 *attS* suicide vector | Cadmium | Dr. Francis Alonzo III (8-9) |
| pJB38 | Shuttle vector with pE194ts temperature sensitive origin for *S. aureus* | Chloramphenicol | BEI Resources |
| pBK123_$P_{cad}$-*recA* | Plasmid with *recA* under the expression of cadmium-inducible promoter ($P_{cad}$) | Chloramphenicol | This study |
| pCM29_$P_{recA}$-*recA-gfp* | Plasmid with *recA* under the expression of its native promoter driving the expression of *gfp* | Chloramphenicol | This study |
| pJC1111_$P_{recA}$-*recA-gfp* | Plasmid with *recA* under its native promoter, translationally fused to *gfp* | Cadmium | This study |
| pJB38_*gyrA*-S84L | pJB38 with *gyrA* containing S84L mutation | Chloramphenicol | This study |
| pJB38_*parC*-S80Y | pJB38 with *parC* containing S80Y mutation | Chloramphenicol | This study |

**Table S3.** *Primers used in this study.*

| Primer | Sequence | Source |
|---|---|---|
| pCM29_*recA*_F | 5′ - TCATCATGCTAGCTCATCGATTGAAAGAAGTGGA - 3′ | This study |
| pCM29_*recA*_R | 5′ - CGATGATGGTACCAGCGAGACCTCCTAATTGAA - 3′ | This study |
| *recA*_pBK123_F | 5′ - CCTGCAGGTCGACTTCAATTAGGAGGTCTC - 3′ | This study |
| *recA*_pBK123_R | 5′ - GCTCGGTACCCGGGGAACTATAGATATAAATTT - 3′ | This study |
| pBK123_seq_F | 5′ - AGAAAAGAAGGAAAACTAGC –3′ | 11 |
| pJC1111_*recA*_Gibson_F | 5′ - CTGAATTCGAGCTCGGTACCTCATCGATTGAAAGAAGTG - 3′ | This study |
| pJC1111_*recA*_Gibson_R | 5′ - GCCTGCAGGTCGACTCTAGACTATTCTTCGTCAAATAATGAC - 3′ | This study |
| pJC1111_*rexB*_Gibson_F | 5′ - CTGAATTCGAGCTCGGTACCTCATCGGTGTTCCTGGTGAT - 3′ | This study |
| pJC1111_*rexB*_Gibson_R | 5′ - GCCTGCAGGTCGACTCTAGACTATTGCTCACCCCCAAATT - 3′ | This study |
| pJC1111_seq_F | 5′ - TGGCCTTTTGCTCACATGTTCTTTCCTGCGTTATCCCCTGATTC - 3′ | 18 |
| pJC1111_seq_R | 5′ - TGATATCAAAATTATACATGTCAACG - 3′ | 18 |
| SaPI_chrom_verify | 5′ - GTGCTTCACCAGCACCACATGCTG - 3′ | 8 |
| SaPI_vector_verify | 5′ - GGTATTAGTTTGAGCTGTCTTGGTTCATTGATTGC - 3′ | 8 |
| SA_*recA*_int_R | 5′ - TCCTTACCTTGACCCATTCG - 3′ | This study |
| *rexB*_Tn_F | 5′ - TCATCGGTGTTCCTGGTGAT - 3′ | This study |
| *xseA*_Tn_F | 5′ - TCACATTACACAAATCGTTATTGAAAA - 3′ | This study |
| Upstream | 5′ - CTCGATTCTATTAACAAGGG - 3′ | 17 |
| Buster | 5′ - GCTTTTTCTAAATGTTTTTTAAGTAAATCAAGTAC - 3′ | 17 |
| pCM29_Gibson_F | 5' - ATGAGCAAAGGAGAAGAACT - 3' | This study |
| pCM29_Gibson_R | 5' - AAATAATCATCCTCCTAAGG - 3' | This study |
| $P_{recA}$_Gibson_F | 5' - CCTTAGGAGGATGATTATTTCATCGATTGAAAGAAGTGGA - 3' | This study |
| $P_{recA}$_Gibson_R | 5' - AGTTCTTCTCCTTTGCTCATTCCTGCGGCGCCTCCTTCTTCGTCAAATAATGACT - 3' | This study |
| pJC1111-*gfp*-R | 5' - GCCTGCAGGTCGACTCTAGAGAATTCTTAGTGGTGGTGGTGGTGGT - 3' | This study |
| *xseA*_F | 5' - AACGTTTGAAGGAGTTAAAAAATTGAGTGATGTTATATGT - 3' | This study |
| *sodA*_F | 5' - ACGTAATAATGGCGGTGGACA - 3′ | This study |
| *katA*_F | 5' - GGGCATCCAGTATCAGACCG - 3′ | This study |
| *tpx*_F | 5' - GGACCAATCCACTTAAAAGGTCA - 3′ | This study |
| pJB38_*gyrA*_F | 5' - CGATGATGAATTCATGGCTGAATTACCTCAATC - 3' | This study |
| pJB38_*gyrA*_R | 5' - CGATGATGGTACCTTATTCTTCATCTGATGATT - 3' | This study |
| pJB38_*parC*_F | 5' - CGATGATGAATTCGCTTGGACAGACGAAGAGCT - 3' | This study |
| pJB38_*parC*_R | 5' - CGATGATGGTACCACCCCTCGCATCCTCTACAT - 3' | This study |
| *parC*_seq | 5' - CAAGTGGTAATACACACGA - 3' | This study |

**Table S4.** *MICs.*

| Compound | WT | *recA* ::Tn | *recA* ::Tn [C] | *rexB* ::Tn | *rexB* ::Tn [C] |
|---|---|---|---|---|---|
| **Delafloxacin (µg/mL)** | 0.002-0.003 | 0.0002-0.0004 | 0.002-0.003 | 0.0002-0.0004 | 0.002-0.003 |
| **Ciprofloxacin (µg/mL)** | 0.008-0.03 | 0.008 | 0.03 | 0.008 | 0.03 |
| **Rifampicin (µg/mL)** | 0.06 | 0.06 | 0.06 | 0.06 | 0.06 |
| **5-Fluoro-2'-Deoxycytidine (µg/mL)** | 0.0002-0.0004 | 0.0001 | 0.0002 | 0.0001 | 0.0002 |
| **2-2'-Bipyridine (mM)** | 1.9-7.5 | 1.9 | 3.8-7.5 | 3.8-7.5 | 3.8-7.5 |
| **Glutathione (mM)** | >7.5 | 1.9-7.5 | 3.8->7.5 | 1.9-7.5 | >7.5 |

**Movie S1 (separate file).** S. aureus *expresses* recA *during nutritive recovery after DLX treatment.* HG003::*$P_{recA}$-gfp* was monitored via timelapse microscopy on a Mueller-Hinton agarose pad after 5 h of treatment with DLX. Green cells are positive for *gfp* expression (indicating *recA* induction), and red cells are positive for PI staining (indicating loss of membrane integrity in dying/dead cells).

**Movie S2 (separate file).** *A second video of* S. aureus *expressing* recA *during nutritive recovery after DLX treatment.*

**Movie S3 (separate file).** S. aureus *expresses* recA *during nutritive recovery after CIP treatment.* HG003::*$P_{recA}$-gfp* was monitored via timelapse microscopy on a Mueller-Hinton agarose pad after 5 h of treatment with CIP. Green cells are positive for *gfp* expression (indicating *recA* induction), and red cells are positive for PI staining (indicating loss of membrane integrity in dying/dead cells).

**Movie S4 (separate file).** *A second video of* S. aureus *expressing* recA *during nutritive recovery after CIP treatment.*

**Movie S5 (separate file).** *Untreated stationary-phase* S. aureus *does not express appreciable levels of* recA *and divides normally.* Untreated HG003::*$P_{recA}$-gfp* was monitored via timelapse microscopy on a Mueller-Hinton agarose pad. No appreciable *gfp* expression (green cells indicating *recA* induction) or PI positivity (red cells indicating loss of membrane integrity) was observed. Additionally, unlike cells recovering from FQ treatment, most untreated cells produce two daughter cells that both give rise to progeny.

**Movie S6 (separate file).** *A second video of untreated* S. aureus *HG003::*$P_{recA}$-gfp *proliferating on a Mueller-Hinton agarose pad.*

**Movie S7 (separate file).** *DLX-treated reporter-free* S. aureus *persisters frequently produce only one successful daughter cell.* HG003 was monitored via timelapse microscopy on a Mueller-Hinton agarose pad after 5 h of treatment with DLX. The persister captured in this video divides into two daughter cells, one of which continues dividing while the other stalls and later stains positive for PI (indicating loss of membrane integrity in dying/dead cells).

**Movie S8 (separate file).** *A second video of DLX-treated reporter-free* S. aureus *HG003 during nutritive recovery.*

**Movie S9 (separate file).** *CIP-treated reporter-free* S. aureus *persisters frequently produce only one successful daughter cell.* HG003 was monitored via timelapse microscopy on a Mueller-Hinton agarose pad after 5 h of treatment with CIP. The persister that is a single cell at the beginning of this video divides into two daughter cells, one of which continues dividing while the other stalls and later stains positive for PI (indicating loss of membrane integrity in dying/dead cells). Groups of cells were labeled "Persister Microcolony" when more than one cell in the group divided, suggesting that these cells arose from a persister that began dividing before our imaging experiment started.

**Movie S10 (separate file).** *A second video of CIP-treated reporter-free* S. aureus *HG003 during nutritive recovery.*

**Movie S11 (separate file).** *Untreated* S. aureus *HG003*::$P_{recA}$-recA-gfp *cells divide normally and form few, faint foci.* Untreated S. aureus *HG003*::*$P_{recA}$-recA-gfp* was monitored via timelapse microscopy on a Mueller-Hinton agarose pad. Cells in later generations occasionally form faint foci that disappear quickly. Some cells in later generations also take up PI (red cells indicating loss of membrane integrity).

**Movie S12 (separate file).** *A second video of untreated* S. aureus *HG003::*$P_{recA}$-recA-gfp *proliferating on a Mueller-Hinton agarose pad.*

**Movie S13 (separate file).** *DLX-treated* S. aureus *HG003*::P$_{recA}$-recA-gfp *persisters and non-persisters form RecA-GFP foci, indicative of DSBs.* HG003::*P$_{recA}$-recA-gfp* cells were treated for 5 h with DLX then monitored during recovery on a Mueller-Hinton agarose pad. Most persisters and non-persisters have green foci that are much brighter than those observed in untreated populations. The persister at the top right forms a RecA-GFP focus before dividing. The group of cells labeled "Persister Microcolony" appears to have arisen from a persister that began dividing before our imaging experiment started. Many cells present at the beginning of recovery lyse and/or turn red, indicating that they have taken up PI.

**Movie S14 (separate file).** *A second video of* S. aureus *HG003::*P$_{recA}$-recA-gfp *during nutritive recovery after DLX treatment.*

**Movie S15 (separate file).** *CIP-treated* S. aureus *HG003*::P$_{recA}$-recA-gfp *persisters and non-persisters form RecA-GFP foci, indicative of DSBs. P$_{recA}$-recA-gfp* cells were treated for 5 h with CIP then monitored during recovery on a Mueller-Hinton agarose pad. Most persisters and non-persisters have green foci that are much brighter than those observed in untreated populations. Two of the three persisters visible in this movie (top right and bottom left) appear to form a RecA-GFP focus before completing cell division. Many cells present at the beginning of recovery lyse and/or turn red, indicating that they have taken up PI.

**Movie S16 (separate file).** *A second video of* S. aureus *HG003::*P$_{recA}$-recA-gfp *during nutritive recovery after CIP treatment.*

**Movie S17 (separate file).** *Post-DLX starvation increases persister fraction and may help persisters divide before forming DSBs. S. aureus* HG003::*P$_{recA}$-recA-gfp* cells were treated for 5 h with DLX, starved for 2 h in PBS, then monitored during recovery on a Mueller-Hinton agarose pad. Unlike unstarved DLX persisters, which usually form RecA-GFP foci before dividing, three of the four persisters visible in this video (all except the bottom one) appear to have no RecA-GFP foci before dividing. Instead, foci first appear in their immediate progeny.

**Movie S18 (separate file).** *A second video of* S. aureus *HG003::*P$_{recA}$-recA-gfp *during nutritive recovery after starvation following DLX treatment.*

**Movie S19 (separate file).** *Post-CIP starvation increases persister fraction and may help persisters divide before forming DSBs. S. aureus* HG003::*P$_{recA}$-recA-gfp* cells were treated for 5 h with CIP, starved for 2 h in PBS, then monitored during recovery on a Mueller-Hinton agarose pad. Unlike unstarved CIP persisters, which usually form RecA-GFP foci before dividing, two of the six persisters visible from the beginning of this video appear to have no RecA-GFP foci before dividing. Instead, foci first appear in their immediate progeny.

**Movie S20 (separate file).** *A second video of* S. aureus *HG003::*P$_{recA}$-recA-gfp *during nutritive recovery after starvation following CIP treatment.*

### *SI* References


1. L. Radlinski, *et al.*, *Pseudomonas aeruginosa* exoproducts determine antibiotic efficacy against *Staphylococcus aureus*. *PLoS Biol* **15**, e2003981 (2017).
2. Seemann, T, Snippy: Rapid bacterial SNP calling and core genome alignments. (2016)
3. P. D. Karp, *et al.*, The BioCyc collection of microbial genomes and metabolic pathways. *Brief Bioinform* **20**, 1085–1093 (2019).
4. S. Fuchs, *et al.*, AureoWiki‐The repository of the *Staphylococcus aureus* research and annotation community. *Int J Med Microbiol* **308**, 558–568 (2018).
5. P. J. Hare, *et al.*, Time-lapse epifluorescence microscopy imaging of *Pseudomonas aeruginosa* and *Staphylococcus aureus* heterogeneous phenotypes. *J Vis Exp* e67617 (2025).
6. K. L. Krausz, J. L. Bose, Bacteriophage transduction in *Staphylococcus aureus*: Broth-based method. *Methods Mol Biol* **1373**, 63–68 (2016).
7. M. E. Olson, Bacteriophage Transduction in *Staphylococcus aureus*. *Methods Mol Biol* **1373**, 69–74 (2016).
8. J. Chen, P. Yoong, G. Ram, V. J. Torres, R. P. Novick, Single-copy vectors for integration at the SaPI1 attachment site for *Staphylococcus aureus*. *Plasmid* **76**, 1–7 (2014).
9. W. P. Teoh, X. Chen, I. Laczkovich, F. Alonzo, *Staphylococcus aureus* adapts to the host nutritional landscape to overcome tissue-specific branched-chain fatty acid requirement. *Proc Natl Acad Sci U S A* **118**, e2022720118 (2021).
10. B. K. Sharma-Kuinkel, *et al.*, The *Staphylococcus aureus* LytSR Two-component regulatory system affects biofilm formation. *J Bacteriol* **191**, 4767–4775 (2009).
11. I. Reslane, *et al.*, Catabolic ornithine carbamoyltransferase activity facilitates growth of *Staphylococcus aureus* in defined medium lacking glucose and arginine. *mBio* **13**, e00395-22 (2022).
12. S. Schenk, R. A. Laddaga, Improved method for electroporation of *Staphylococcus aureus*. *FEMS Microbiol Lett* **94**, 133–138 (1992).
13. M. R. Grosser, A. R. Richardson, Method for preparation and electroporation of *S. aureus* and *S. epidermidis*. *Methods Mol Biol* **1373**, 51–57 (2016).
14. B. N. Kreiswirth, *et al.*, The toxic shock syndrome exotoxin structural gene is not detectably transmitted by a prophage. *Nature* **305**, 709–712 (1983).
15. Y. Y. Pang, *et al.*, *agr*-dependent interactions of *Staphylococcus aureus* USA300 with human polymorphonuclear neutrophils. *J Innate Immun* **2**, 546–559 (2010).
16. H. Veiga, A. M. Jorge, M. G. Pinho, Absence of nucleoid occlusion effector Noc impairs formation of orthogonal FtsZ rings during *Staphylococcus aureus* cell division. *Molec Microbiol* **80**, 1366–1380 (2011).
17. P. D. Fey, *et al.*, A genetic resource for rapid and comprehensive phenotype screening of nonessential *Staphylococcus aureus* genes. *mBio* **4**, e00537-12 (2013).
18. M. Podkowik, *et al.*, Quorum-sensing *agr* system of *Staphylococcus aureus* primes gene expression for protection from lethal oxidative stress. *eLife* **12** (2023).
19. J. L. Bose, P. D. Fey, K. W. Bayles, Genetic Tools To Enhance the Study of Gene Function and Regulation in Staphylococcus aureus. *Appl Environ Microbiol* **79**, 2218–2224 (2013).
20. J. M. Remy, C. A. Tow-Keogh, T. S. McConnell, J. M. Dalton, J. A. Devito, Activity of delafloxacin against methicillin-resistant *Staphylococcus aureus*: Resistance selection and characterization. *J Antimicrob Chemother* **67**, 2814–2820 (2012).
21. Clinical and Laboratory Standards Institute, Performance standards for antimicrobial susceptibility testing. 31st ed. (2021).
22. J. M. Lensmire, *et al.*, The glutathione import system satisfies the *Staphylococcus aureus* nutrient sulfur requirement and promotes interspecies competition. *PLoS Genet* **19**, e1010834 (2023).
23. M. Goswami, N. Jawali, N-acetylcysteine-mediated modulation of bacterial antibiotic susceptibility. *Antimicrob Agents Chemother* **54**, 3529–3530 (2010).
24. F. Wong, *et al.*, Reactive metabolic byproducts contribute to antibiotic lethality under anaerobic conditions. *Mol Cell* **82**, 3499-3512.e10 (2022).

25. A. A. Potter, D. R. Musgrave, J. S. Loutit, Thymine Metabolism in *Pseudomonas aeruginosa* Strain 1: The Presence of a Salvage Pathway. *Microbiology* **128**, 1391–1400 (1982).
26. P. J. Hare, J. R. Gonzalez, R. M. Quelle, Y. I. Wu, W. W. K. Mok, Metabolic and transcriptional activities underlie stationary-phase *Pseudomonas aeruginosa* sensitivity to Levofloxacin. *Microbiol Spectr* **12**, e03567-23.
27. J. I. Batchelder, A. J. Taylor, W. W. K. Mok, Metabolites augment oxidative stress to sensitize antibiotic-tolerant *Staphylococcus aureus* to fluoroquinolones. *mBio* **15**, e02714-24 (2024).
28. A. Ducret, E. M. Quardokus, Y. V. Brun, MicrobeJ, a tool for high throughput bacterial cell detection and quantitative analysis. *Nat Microbiol* **1**, 16077 (2016).
29. D. Ershov, *et al.*, TrackMate 7: Integrating state-of-the-art segmentation algorithms into tracking pipelines. *Nat Methods* **19**, 829–832 (2022).
30. J.-Y. Tinevez, *et al.*, TrackMate: An open and extensible platform for single-particle tracking. *Methods* **115**, 80–90 (2017).
31. T. J. LaGree, B. A. Byrd, R. M. Quelle, S. L. Schofield, W. W. K. Mok, Stimulating transcription in antibiotic-tolerant *Escherichia coli* sensitizes it to fluoroquinolone and nonfluoroquinolone topoisomerase inhibitors. *Antimicrob Agents Chemother* **67**, e0163922 (2023).
